# Interstitial macrophages drive chronic lung allograft dysfunction

**DOI:** 10.64898/2026.08.21.746267

**Authors:** Atsushi Suzuki, Maxwell J. Schleck, Qiang Wu, Radmila A. Fenton, Luisa Cusick, Taisuke Kaiho, Hiam Abdala-Valencia, Zhan Yu, Yuliana V. Sokolenko, Ziyan Lu, Suchitra Swaminathan, Mary Carns, Suror Mohsin, Phillip Cooper, Vikas Mehta, Taichi Nagano, Lee A.D. Cooper, Mrinalini Venkata Subramani, Catherine N. Myers, Ambalavanan Arunachalam, Chitaru Kurihara, Ankit Bharat, G.R. Scott Budinger, Alexander V. Misharin

## Abstract

Despite immunosuppressive regimens targeting adaptive immunity, chronic lung allograft dysfunction (CLAD) remains the major obstacle to durable lung allograft survival. Here, we identify colony-stimulating factor 1 receptor (CSF1R)-expressing interstitial macrophages as critical orchestrators of CLAD. Using lung tissue from patients with CLAD and a mouse model of mismatched lung transplantation, we show that both donor-derived tissue-resident and recipient-monocyte-derived interstitial macrophages spatially co-localize within peribronchial immune aggregates in patients with CLAD. These interstitial macrophages express distinct cytokine programs that include those implicated in the recruitment of T and B cells. Pharmacological inhibition of CSF1R after lung transplantation in mice reduced interstitial macrophage abundance and attenuated CLAD pathology. Our findings identify donor- and recipient-derived interstitial macrophages as upstream regulators of CLAD and suggest CSF1R as a therapeutic target for its prevention and treatment.

## INTRODUCTION

Lung transplantation is a life-saving intervention for patients with advanced lung diseases, but long-term survival is limited by rejection of the allograft, recognized as the clinical syndrome chronic lung allograft dysfunction (CLAD). Despite improvements in perioperative management and chronic immunosuppression, CLAD develops in 40–50% of recipients within five years and is the major barrier to long-term survival after lung transplantation.^1,2^

Bronchiolitis obliterans syndrome (CLAD-BOS), the most common presentation of CLAD, is characterized by airflow obstruction resulting from fibroblast proliferation and activation around small airways accompanied by the peribronchial accumulation of lymphoid infiltrates.^3–7^ Persistent immune activation in response to donor antigens, the expansion of donor-reactive T and B cell clones, and the formation of tertiary lymphoid structures (TLS) have been implicated in the pathogenesis of CLAD-BOS.^8^ Nevertheless, immunosuppressive therapies targeting lymphocytes have limited efficacy, at best modestly slowing the progression of CLAD-BOS without reversing it.^9,10^ These observations suggest that additional cellular populations sustain the organization of lymphoid structures and the associated fibroproliferation in the lung allograft.

Macrophages are observed within the peribronchial lymphocytic infiltrates in patients with CLAD-BOS, but their ontogeny, spatial organization, and role in its pathogenesis are unclear.^7,11,12^ This question takes on clinical relevance with the demonstration that an inhibitory antibody targeting the CSF1R, axatilimab, reversed established lung fibrosis in the lungs of patients with chronic graft-versus-host-disease (cGVHD) after hematopoietic stem cell transplantation, a disorder that shares some histologic features with CLAD-BOS.^7,13^ Because CSF1R expression is largely restricted to monocytes, macrophages, and dendritic cells, and tonic CSF1R signaling is required for monocyte and macrophage maintenance,^14–16^ the improvement in lung fibrosis observed with axatilimab implicates these populations in sustaining fibrotic airway disease.

Here, we used lineage-tracing and single-cell spatial transcriptomic analyses of human lung explants from patients with CLAD-BOS and a murine model of CLAD-BOS after lung transplantation to identify interstitial macrophages as organizers of the peribronchial lymphoid niche characteristic of CLAD-BOS. These populations originated from the expansion of donor-derived tissue-resident interstitial macrophages and the recruitment of recipient monocytes that differentiated into monocyte-derived interstitial macrophages. Although donor- and recipient-derived interstitial macrophages expressed distinct cytokine programs, both converged on the recruitment and activation of T and B cells. In mice, CSF1R inhibition depleted both interstitial macrophage populations, and prevented the accumulation of T and B cells in peribronchial lymphoid aggregates, mitigating CLAD-BOS pathology. Our findings suggest that interstitial macrophages originating from the donor and recipient orchestrate the organization of innate immune cells around the airways to promote CLAD-BOS. Targeting CSF1R signaling in interstitial macrophages could represent a novel therapeutic strategy for these patients.

## RESULTS

### Interstitial macrophages are expanded in patients with CLAD-BOS

We performed single-cell spatial transcriptomic analysis of explanted lung tissues from patients with CLAD-BOS (n = 3). As normal lung controls, we included surgical lung biopsies of donor lungs obtained at the time of lung transplantation (n = 4). As fibrotic lung controls, we included biopsies of lung explants from patients with idiopathic pulmonary fibrosis (IPF). We chose IPF because it is characterized by progressive fibrosis in the alveolar space with relative sparing of the airways. In contrast, CLAD-BOS is characterized by fibrosis around small airways in the bronchovascular bundle, a confined anatomical space proximal to the alveolar region where bronchi and blood vessels are enclosed in a connective tissue sheath (**Suppl. Figure S1a, b**). For all samples, we analyzed a large contiguous tissue area encompassing the bronchovascular bundle, alveolar region, and pleural surface (**Figure 1a, Suppl. Figure S2**).

**Figure 1.**
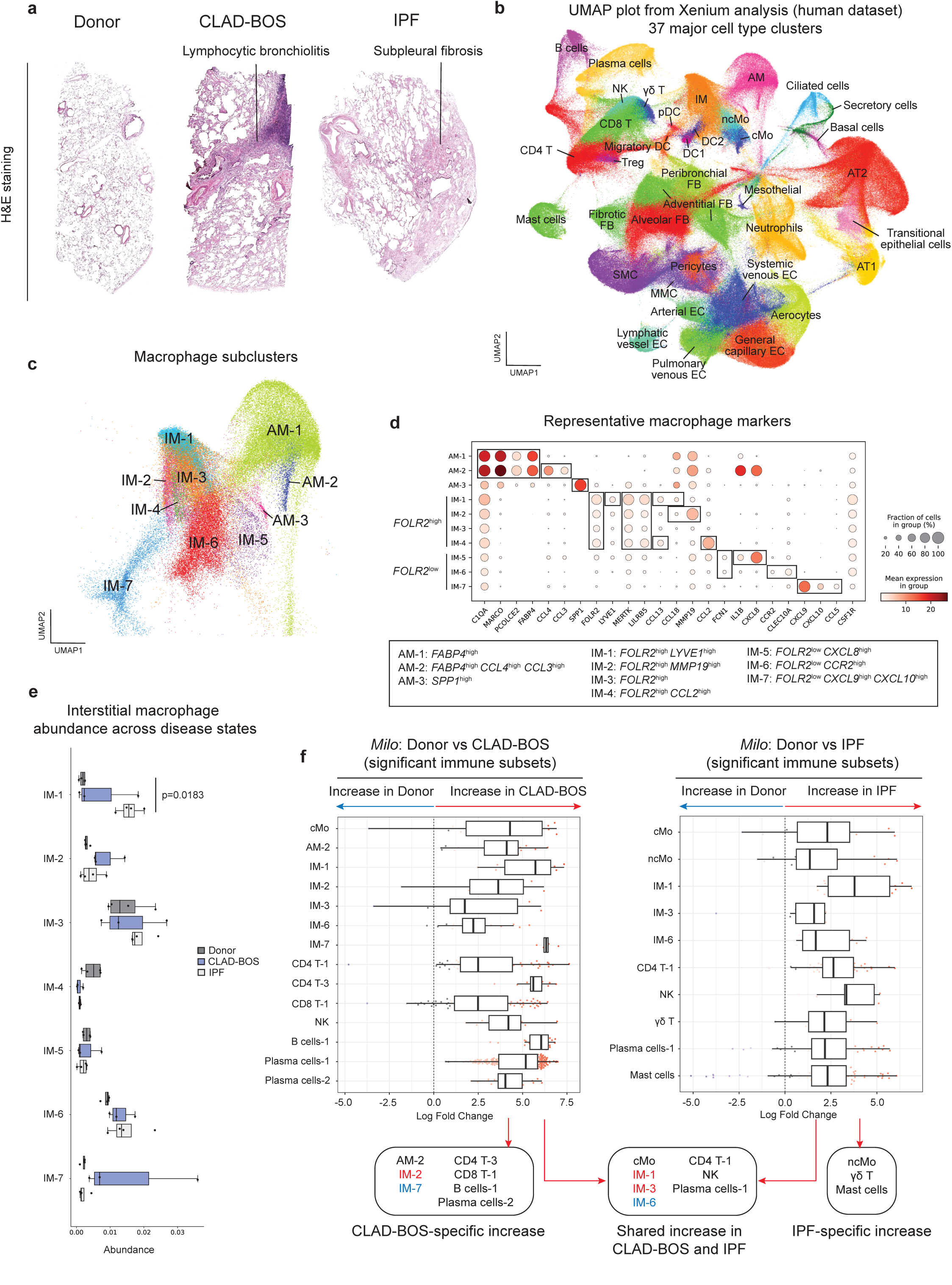
Interstitial macrophages are expanded in patients with CLAD-BOS. **a**. Representative hematoxylin and eosin (H&E)-stained images of the donor lung (n = 4), and lungs from patients with CLAD-BOS (n = 3) and IPF (n = 4) analyzed using the Xenium platform. These samples included the bronchovascular bundle, alveolar region, and pleural surface. **b.** Uniform manifold approximation and projection (UMAP) visualization of single-cell spatial transcriptomic data from human samples (total number of cells = 881,761 cells, 389 genes, 37 major cell type clusters). AM = alveolar macrophages; AT1 = alveolar epithelial type I cells; AT2 = alveolar epithelial type II cells; CD4 T = CD4 T cells; CD8 T = CD8 T cells; cMo = classical monocytes; DC = dendritic cells; EC = endothelial cells; FB = fibroblasts; γδ T = gamma delta T cells; IM = interstitial macrophages; MMC = microvascular mural cells; ncMo = non-classical monocytes; NK = natural killer cells; pDC = plasmacytoid dendritic cells; SMC = smooth muscle cells; Treg = regulatory T cells. **c.** UMAP visualization of macrophage subsets. **d.** Dot plot showing expression of genes distinguishing each macrophage subtype. *FOLR2*^high^ interstitial macrophages = IM-1-4; *FOLR2*^low^ interstitial macrophages = IM-5-7. **e.** Differential abundance analysis of interstitial macrophage subsets based on per-sample fractions. Box plots represent the median and interquartile range. Statistical significance was assessed using pairwise *t*-tests with Bonferroni correction for multiple comparisons. **f.** Differential abundance analysis using *Milo* showing significantly enriched immune cell subsets in CLAD-BOS and IPF compared to donor lungs. Box plots represent the median and interquartile range. Cell populations were defined as significantly changed if at least four neighborhoods were identified, and both the upper and lower quartiles of log-fold change were skewed to one side relative to zero.

We identified 37 major cell types (**Figure 1b**), which were further resolved into 62 transcriptionally distinct subclusters (**Suppl. Figure S3a-j**). Based on transcriptional signatures and spatial distributions, we resolved alveolar and interstitial macrophages (**Figure 1c, d**). Within alveolar macrophages, we resolved two populations of *FABP4^high^*tissue-resident alveolar macrophages: *CCL4*^low^*CCL3*^low^*FABP4*^high^ (AM-1) and *CCL4*^high^*CCL3*^high^*FABP4*^high^ (AM-2). We also resolved *SPP1*^high^ monocyte-derived alveolar macrophages (AM- 3).^17–19^ Within the interstitial macrophages, we resolved seven transcriptionally distinct subsets characterized by distinct cytokine and chemokine programs. Using nomenclature defined by others,^20,21^ we grouped interstitial macrophages into *FOLR2*^high^ (IM-1-4) and *FOLR2*^low^ (IM-5-7) subsets based on their *FOLR2* expression (**Figure 1d**). All interstitial macrophage subsets were detected in every sample (**Suppl. Figure S4a**).

Differential abundance analysis of per-sample cell fractions demonstrated compositional changes across many cell types in CLAD-BOS and IPF (**Figure 1e, Suppl. Figure S4b**). Differential abundance testing using *Milo* ^22^ revealed both shared and disease-specific changes in cell type abundance between CLAD-BOS and IPF in comparison to donor lungs (**Figure 1f**, **Suppl. Figure S5a, b**). Both CLAD-BOS and IPF lungs showed an increased proportion of *FOLR2*^high^ interstitial macrophages (IM-1, IM-3), *FOLR2*^low^ interstitial macrophages (IM-6), classical monocytes, NK cells, and specific subsets of CD4 T cells, and plasma cells. In contrast, distinct immune subsets were restricted to a single condition. Specifically, *FOLR2*^high^ *MMP19*^high^ interstitial macrophages (IM-3), *FOLR2*^low^ *CXCL9*^high^ *CXCL10*^high^ interstitial macrophages (IM-7), *CXCL13*^high^ CD4 T cells (CD4 T-3), CD8 T cells, B cells, and *IRF4*^low^ plasma cells (Plasma cells-2) were enriched in CLAD-BOS, whereas non-classical monocytes, γδ T cells, and mast cells were enriched in IPF. Together, this analysis identified shared and disease-specific changes in lung tissue composition in patients with CLAD-BOS and IPF.

### *FOLR2*^high^ interstitial macrophages are the predominant macrophage subset in perivascular and alveolar stromal niches in donor lungs

Next, using *CellCharter*^23^, we identified 9 multicellular niches corresponding to known microanatomical structures annotated by a pulmonary pathologist. These spatial domains comprised the perivascular immune niche, peribronchial TLS niche, airspace niche, airway epithelial-fibroblast niche, lymphatic-fibroblast niche, alveolar epithelial niche, alveolar capillary niche, alveolar stromal niche, and arterial endothelial niche (**Figure 2a**, **Suppl. Figure S6a, b**). The cellular composition of these niches revealed highly specialized microenvironments. The perivascular immune niche was characterized by the accumulation of various immune cells, including interstitial macrophage subsets, CD4 and CD8 T cells, NK cells, dendritic cells (DC), and mast cells. The peribronchial TLS niche shared components with the perivascular immune niche, including interstitial macrophages and lymphocytes, but was distinguished by the accumulation of B cells and plasma cells. The similarity between these two niches was confirmed by hierarchical clustering of niche composition correlations (**Figure 2b**). The airspace niche contained predominantly alveolar macrophages. The lymphatic-fibroblast niche, consisting of lymphatic endothelial cells and adventitial fibroblasts, was localized to the perivascular adventitia and pleura of donor lungs (**Figure 2f**). The airway epithelial-fibroblast, alveolar epithelial, and arterial endothelial niches represented corresponding structural compartments of the lung parenchyma. The alveolar capillary niche contained general capillary endothelial cells and aerocytes, whereas the alveolar stromal niche contained pericytes, alveolar fibroblasts, and interstitial macrophage subsets. Notably, quantitative evaluation of niche abundance revealed disease-specific differences (**Figure 2c**). The peribronchial TLS niche, which was virtually absent in donor lungs, was significantly expanded in CLAD-BOS, whereas the perivascular immune, airway epithelial-fibroblast, and lymphatic-fibroblast niches were expanded in IPF.

**Figure 2.**
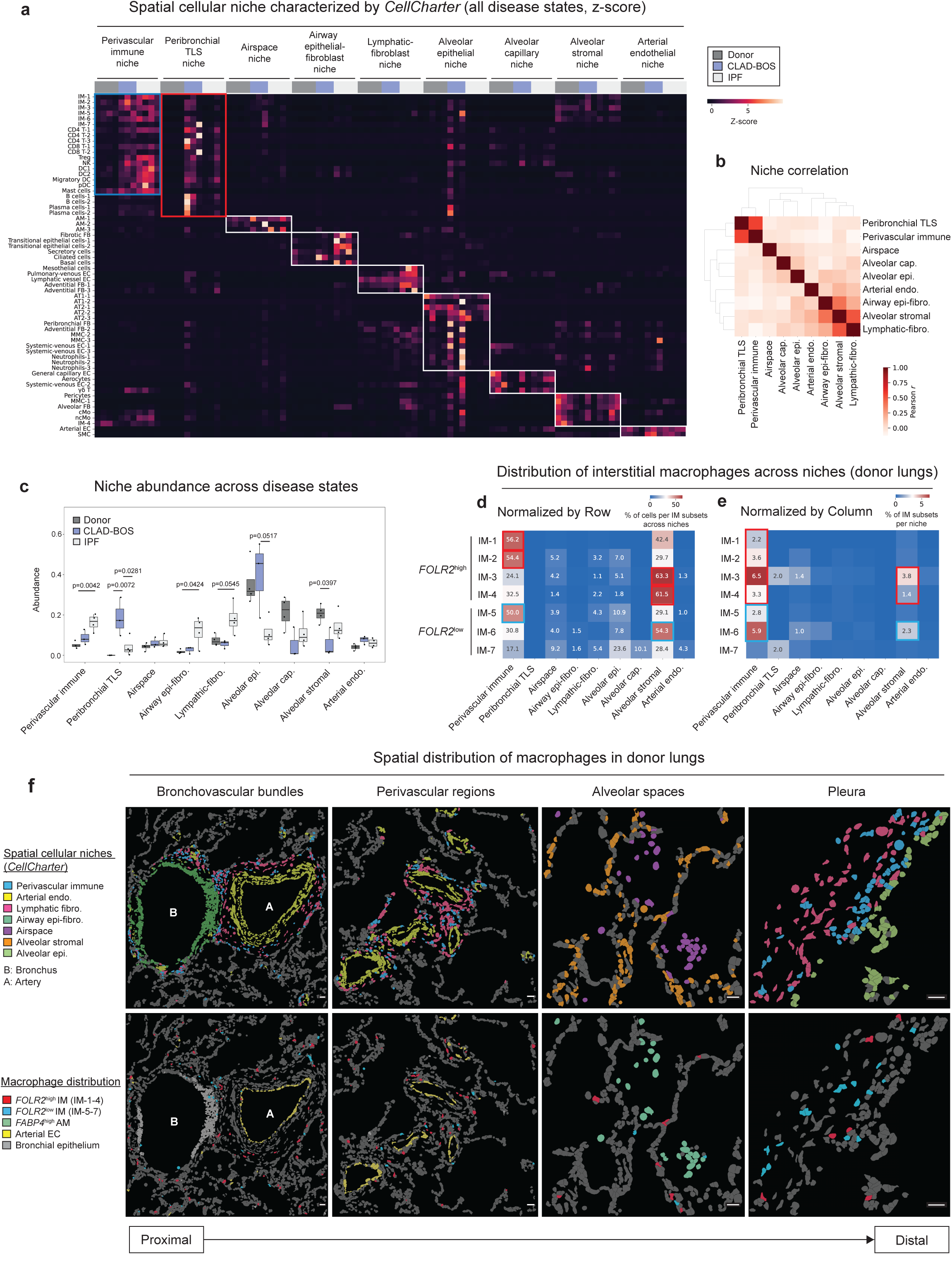
*FOLR2*^high^ interstitial macrophages are the predominant macrophage subset in perivascular and alveolar stromal niches in donor lungs. **a.** Heatmap illustrating the enrichment of specific cell populations within each niche (z-score). **b.** Pearson correlation matrix showing similarity between niches based on their composition. **c.** Comparison of niche abundance across disease groups. Box plots represent the median and interquartile range. Statistical significance was assessed using pairwise *t*-tests with Bonferroni correction for multiple comparisons. **d.** Distribution of interstitial macrophages across niches in donor lungs (normalized by row). **e.** Distribution of interstitial macrophages across niches in donor lungs (normalized by column). The heatmap displays the proportion of cells in each interstitial macrophage subset across niches, calculated by pooling cells from all samples. Values representing less than 1% are not shown. **f.** Representative images showing spatial cellular niches and the distribution of macrophage subsets across the proximal-to-distal axis of donor lungs. Selected cell types are highlighted in each panel. Scale bars 25 μm.

Next, we characterized the spatial distribution of interstitial macrophages across niches. In donor lungs, interstitial macrophages were predominantly localized to the perivascular immune and alveolar stromal niches (**Figure 2d-f**). Within these two niches, the *FOLR2*^high^ subset (IM-1-4) represented the majority of interstitial macrophages (perivascular immune niche: *FOLR2*^high^ 62.6% vs *FOLR2*^low^ 37.4%; alveolar stromal niche: *FOLR2*^high^ 67.3% vs *FOLR2*^low^ 32.6%), consistent with their distribution in the spatial mapping (**Figure 2f**). Together, these findings localize interstitial macrophages to distinct pathological niches in CLAD-BOS and IPF.

### *FOLR2*^high^ and *FOLR2*^low^ interstitial macrophages co-localize with lymphocytes in CLAD-BOS-specific niches

We further examined the spatial localization of *FOLR2*^high^ and *FOLR2*^low^ interstitial macrophages in samples from patients with CLAD-BOS compared to those with IPF. While few *FOLR2*^high^ and *FOLR2*^low^ interstitial macrophages were present in the donor lung, they were present within the expanded CLAD-BOS-specific peribronchial TLS niche (**Figure 2c, 3a–c**). The pathologic correlate of CLAD-BOS is obliterative bronchiolitis, a patchy fibro-inflammatory process circumferentially surrounding the small airways, including the respiratory bronchioles. This pathology was localized to the airway epithelial-fibroblast niche, where fibrotic fibroblasts expressing *CTHRC1*, also seen in samples from patients with IPF, resided (**Figure 3d, Suppl. Figure S1a, S5a, b**). The pathology extended outward from the airways to the peribronchial TLS niche, surrounded by the lymphatic-fibroblast niche and perivascular immune niches, and finally to the adjacent vessel in the bronchovascular bundle identified by the arterial endothelial niche (**Figure 3d, e).** Differential abundance testing using *Milo* to compare niche composition revealed that the peribronchial TLS niche is enriched for multiple lymphocyte populations, including CD4 and CD8 T cells, B cells, and plasma cells, while the perivascular immune niche was characterized by an enrichment of NK cells and mast cells (**Figure 3f**).

**Figure 3.**
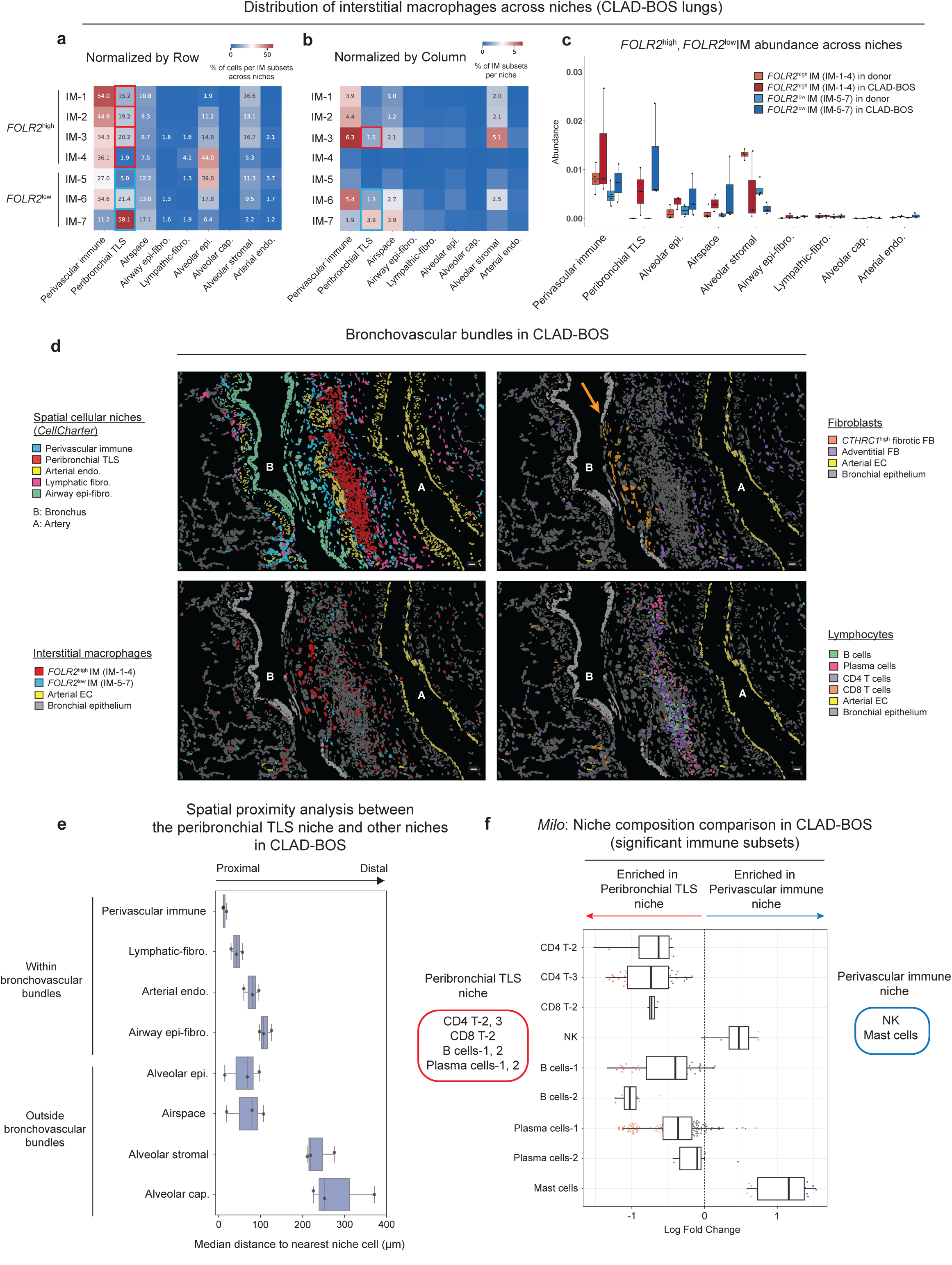
*FOLR2*^high^ and *FOLR2*^low^ interstitial macrophages co-localize with lymphocytes in CLAD-BOS-specific niches. **a.** Distribution of interstitial macrophages across niches in CLAD-BOS lungs (normalized by row). **b.** Distribution of interstitial macrophages across niches in CLAD-BOS lungs (normalized by column). The heatmap displays the proportion of cells in each interstitial macrophage subset across niches, calculated by pooling cells from all samples. Values representing less than 1% are not shown. **c.** Relative abundance of *FOLR2*^high^ and *FOLR2*^low^ interstitial macrophages across niches, comparing donor vs. CLAD-BOS. Box plots represent the median and interquartile range. **d.** Representative images highlighting the distribution of spatial cellular niches and their constituent cell types within the bronchovascular bundle of CLAD-BOS lungs. Arrow indicates the accumulation of *CTHRC1*^high^ fibrotic fibroblasts (FB) around the bronchi. Selected cell types are highlighted in each panel. Scale bars 25 μm. **e.** Spatial proximity analysis between the peribronchial TLS niche and other niches in CLAD-BOS lungs. Box plots showing the median absolute distance (μm) from edge cells within the peribronchial TLS niche to the nearest cells of other niches (n = 3). **f.** Differential abundance analysis using *Milo* showing immune cell types significantly enriched in the peribronchial TLS niche compared to the perivascular immune niche. All box plots represent the median and interquartile range.

Findings in patients with IPF significantly differed from those in patients with CLAD-BOS. Specifically, in the lungs from patients with IPF, interstitial macrophages, particularly the *FOLR2*^high^ subsets (IM-1-4), accumulated in the perivascular immune niche (**Suppl. Figure S7a-c**). Mapping single-cell spatial transcriptomic data onto pathologist-annotated regions showed that fibroblastic foci were characterized by the airway epithelial-fibroblast niche, consisting of *CDH2*^high^ transitional epithelial cells and *CTHRC1*^high^ fibrotic fibroblasts, while advanced fibrotic regions contained the lymphatic fibroblast niche (**Suppl. Figure S7d, e**). Interestingly, the perivascular immune niche enriched for *FOLR2*^high^ interstitial macrophages was found exclusively in advanced fibrotic regions and was absent from fibroblastic foci (**Suppl. Figure S7f**). This was also supported by proximity analysis, confirming the close spatial proximity between the perivascular immune niche and the lymphatic fibroblast niche (**Suppl. Figure S7g**). Next, we analyzed the cellular composition of the perivascular immune niche and identified an enrichment for mast cells (**Suppl. Figure S7h**). Within this niche, mast cells were spatially localized adjacent to *FOLR2*^high^ interstitial macrophages and expressed *CSF1* (**Suppl. Figure S7i, j**).

Given that peribronchial TLS niches were specific to CLAD-BOS, these findings support our hypothesis that interstitial macrophages and lymphocytes may crosstalk within the niche and synergistically contribute to the pathogenesis of CLAD-BOS.

### *FOLR2*^high^ and *FOLR2*^low^ interstitial macrophages display distinct immunoregulatory transcriptomic profiles in CLAD-BOS

To infer the function of interstitial macrophage subsets in CLAD-BOS, we performed a re-analysis of three publicly available single-cell RNA-seq (scRNA-seq) datasets encompassing lung samples from patients with CLAD-BOS and donors (GSE224210, GSE290834, and GSE289881; total: CLAD-BOS, n = 13; Donor, n = 11) (**Suppl. Figure S8a-S8d**). Consistent with our spatial transcriptomic analysis, integrative analysis of scRNA-seq data identified clusters of *FOLR2*^high^ (scIM-1, 2) and *FOLR2*^low^ interstitial macrophages (scIM-3-5) (**Figure 4a**). We focused on the *FOLR2*^high^ scIM-2 and *FOLR2*^low^ scIM-4 clusters, which were represented across all datasets (**Suppl. Figure S8e**). The representative markers distinguishing *FOLR2*^high^ and *FOLR2*^low^ interstitial macrophages recapitulated the transcriptomic profiles observed in our spatial analysis (**Figure 4b**). Briefly, *FOLR2*^high^ interstitial macrophages (scIM-2) expressed *CCL18*, *CCL13*, and *CCL2*, which are potent chemoattractants for monocytes and T cells, whereas *FOLR2*^low^ interstitial macrophages (scIM-4) expressed *CXCL9* and *CXCL10*, key recruiters of CXCR3-expressing CD4 and CD8 T cells. Next, we identified differentially expressed genes (DEGs) between donor lungs and lungs from patients with CLAD-BOS in *FOLR2*^high^ and *FOLR2*^low^ subsets (**Figure 4c, Suppl. Table S4**). *FOLR2*^high^ interstitial macrophages (scIM-2) in CLAD-BOS upregulated the stress-induced cytokine *GDF15*, the lymphocyte-checkpoint ligand *PDCD1LG2* (PD-L2), and the retinoic acid synthase *RDH10*, among others. In contrast, *FOLR2*^low^ interstitial macrophages (scIM-4) exhibited upregulation of the T-cell chemoattractants (*CXCL9* and *CXCL10*), as well as the T cell costimulatory molecule *CD80*. To characterize the functional differences between these subsets in the context of CLAD-BOS, we performed gene set enrichment analysis (GSEA) using 50 hallmark gene sets from MSigDB (**Figure 4d**). *FOLR2*^high^ interstitial macrophages were enriched in gene sets related to tissue metabolic adaptation and homeostasis, including “XENOBIOTIC METABOLISM” (*EPHX1*, *APOE*, *RBP4*, and *CYP27A1*), “ADIPOGENESIS” (*APOE*, *LPL*, and *ME1*), and “COAGULATION” (*GNG12* and *DPP4*). In contrast, *FOLR2*^low^ interstitial macrophages were enriched in gene sets related to inflammatory and immune-response pathways, including “ALLOGRAFT REJECTION” (*CCL22*, *CD1D*, *CCR2*, and *EREG*), “INFLAMMATORY RESPONSE” (*OSM*, *CLEC5A*, *CCL22*, *MEFV*, and *EREG*), and “INTERFERON GAMMA RESPONSE” (*IDO1*). Overall, these analyses demonstrated that in patients with CLAD-BOS, *FOLR2*^high^ interstitial macrophages exhibit immunoregulatory profiles distinct from *FOLR2*^low^ interstitial macrophages.

**Figure 4.**
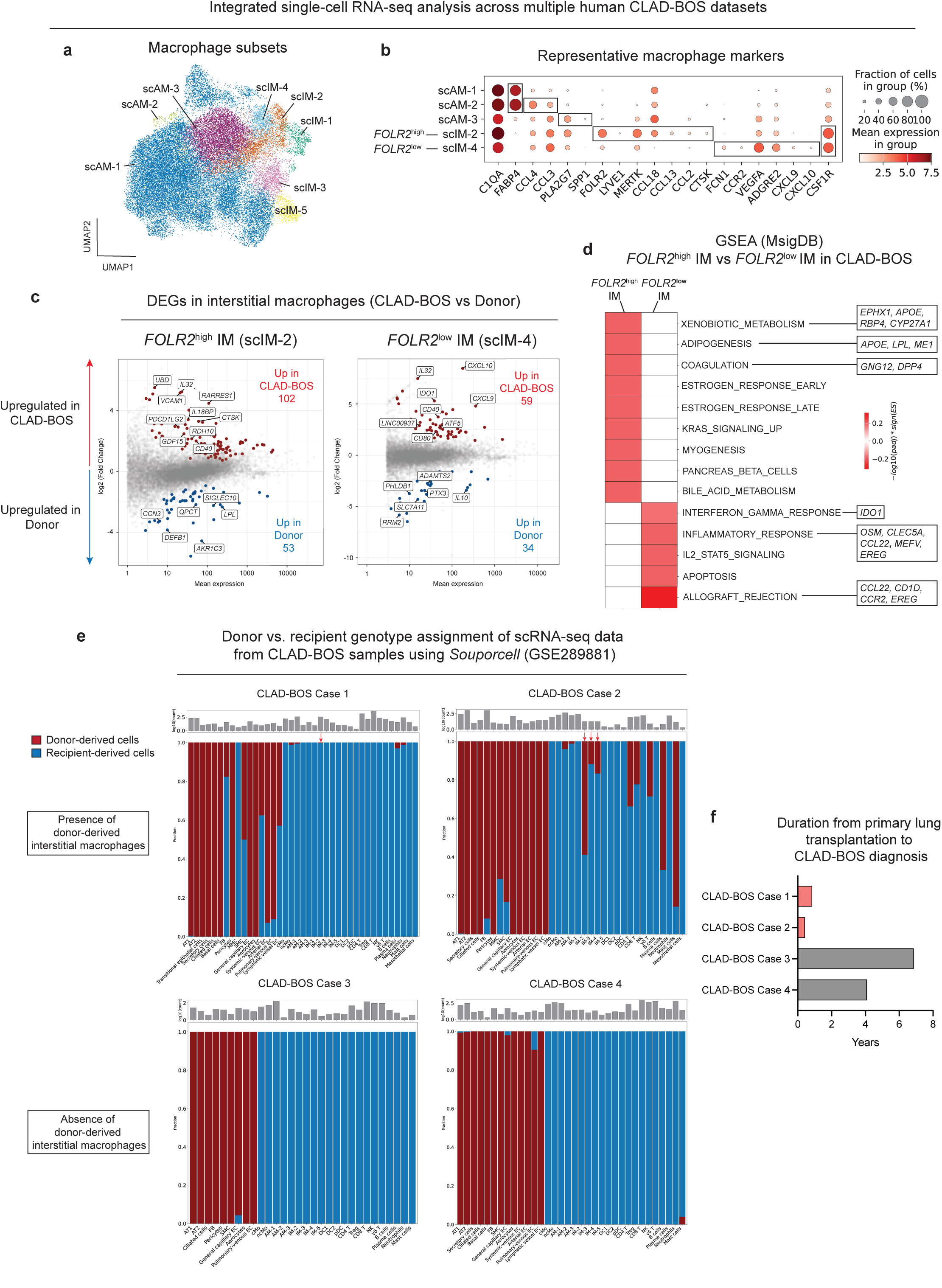
*FOLR2*^high^ and *FOLR2*^low^ interstitial macrophages display distinct immunoregulatory transcriptomic profiles in CLAD-BOS. **a.** UMAP visualization of macrophage subsets from the integrated analysis of three independent, publicly available CLAD-BOS scRNA-seq datasets (GSE224210, GSE2908334, GSE289881; total: CLAD-BOS, n = 13; Donor, n = 11). A UMAP visualization of all cell types is provided in Suppl. Figure S8. **b.** Dot plot showing expression of genes distinguishing each macrophage subtype. **c.** Mean-Average (MA) plots displaying differentially expressed genes (DEGs) in *FOLR2*^high^ interstitial macrophages (scIM-2) and *FOLR2*^low^ interstitial macrophages (scIM-4) when comparing CLAD-BOS versus donor lungs. DEGs with a |log_2_(fold-change)| > 0.585 (representing a ≥ 1.5-fold change in expression) and an adjusted *p*-value < 0.05 are plotted. The key representative markers are highlighted. **d.** Heatmap showing the results of Gene Set Enrichment Analysis (GSEA) using 50 hallmark gene sets from MsigDB, comparing *FOLR2*^high^ vs *FOLR2*^low^ interstitial macrophages in CLAD-BOS. Enrichment scores (ES) and adjusted p-values (*padj*) are represented by the value -log_10_ (*padj*) X sign (*ES*). The heatmap displays pathways for which the absolute value of -log_10_ (*padj*) X sign (*ES*) exceeds 0.15. Representative genes associated with specific gene sets are highlighted in the boxes on the right. **e.** Donor vs. recipient genotype assignment of scRNA-seq data from CLAD-BOS samples using *Souporcell* (n = 4). **f.** Duration from primary lung transplantation to CLAD-BOS diagnosis for samples included in *Souporcell* analysis (n = 4).

### Recipient monocyte-derived interstitial macrophages predominate in human CLAD-BOS

Given the unique context of transplantation, where donor- and recipient-derived cells can coexist as an acquired chimera, we explored whether donor-derived interstitial macrophages persist long-term post-transplant or are replaced by recipient monocyte-derived interstitial macrophages. To test this, we utilized the *Souporcell* pipeline^24^ to perform genotype-based deconvolution of a scRNA-seq dataset for which raw FASTQ files were available (n = 4, GSE289881). This allowed us to assign donor- or recipient-origin to individual cells. This analysis demonstrated that while donor-derived interstitial macrophages persisted and were detectable in patients with CLAD-BOS at the time of re-transplantation up to 16 months post-transplant, recipient-derived interstitial macrophages were predominant (**Figure 4e**). Interestingly, the two patients with detectable donor-derived interstitial macrophages were diagnosed with CLAD-BOS considerably earlier than the two patients in whom these cells were absent (0.4- and 0.9-years vs 4.1- and 6.9-years, respectively; **Figure 4f**). Together, these findings indicate that both donor- and recipient-derived interstitial macrophages could contribute to the pathogenesis of CLAD-BOS.

### CSF1R blockade ameliorates CLAD-BOS pathology

Therapeutic blockade of CSF1R signaling with axatilimab demonstrated efficacy in patients with cGVHD, which shares some histopathologic features with CLAD-BOS.^7,13^ Thus, to further investigate the ontogeny and causal role of interstitial macrophages in the pathogenesis of CLAD-BOS, we blocked CSF1R using the PLX3397 (Pexidartinib) in a murine model of CLAD-BOS based on orthotopic allogeneic minor antigen-mismatched lung transplantation (*HLA-A2.1* knock-in donor lungs into C57BL/6J recipients)^3,4^ (**Figure 5a**). This model recapitulates key biological features of human CLAD-BOS, including T and B cell responses against donor tissues and BOS-like pathology.^3,4^ Syngeneic lung transplant recipients (C57BL/6J donor lungs into C57BL/6J recipients) served as controls (**Figure 5a**). As expected, the allogeneic lung transplant resulted in the development of BOS-like pathology, characterized by inflammatory cell infiltrates along the bronchovascular bundles (**Figure 5b, 5c)**. Treatment with PLX3397 ameliorated this pathology (**Figure 5b-5d).**

**Figure 5.**
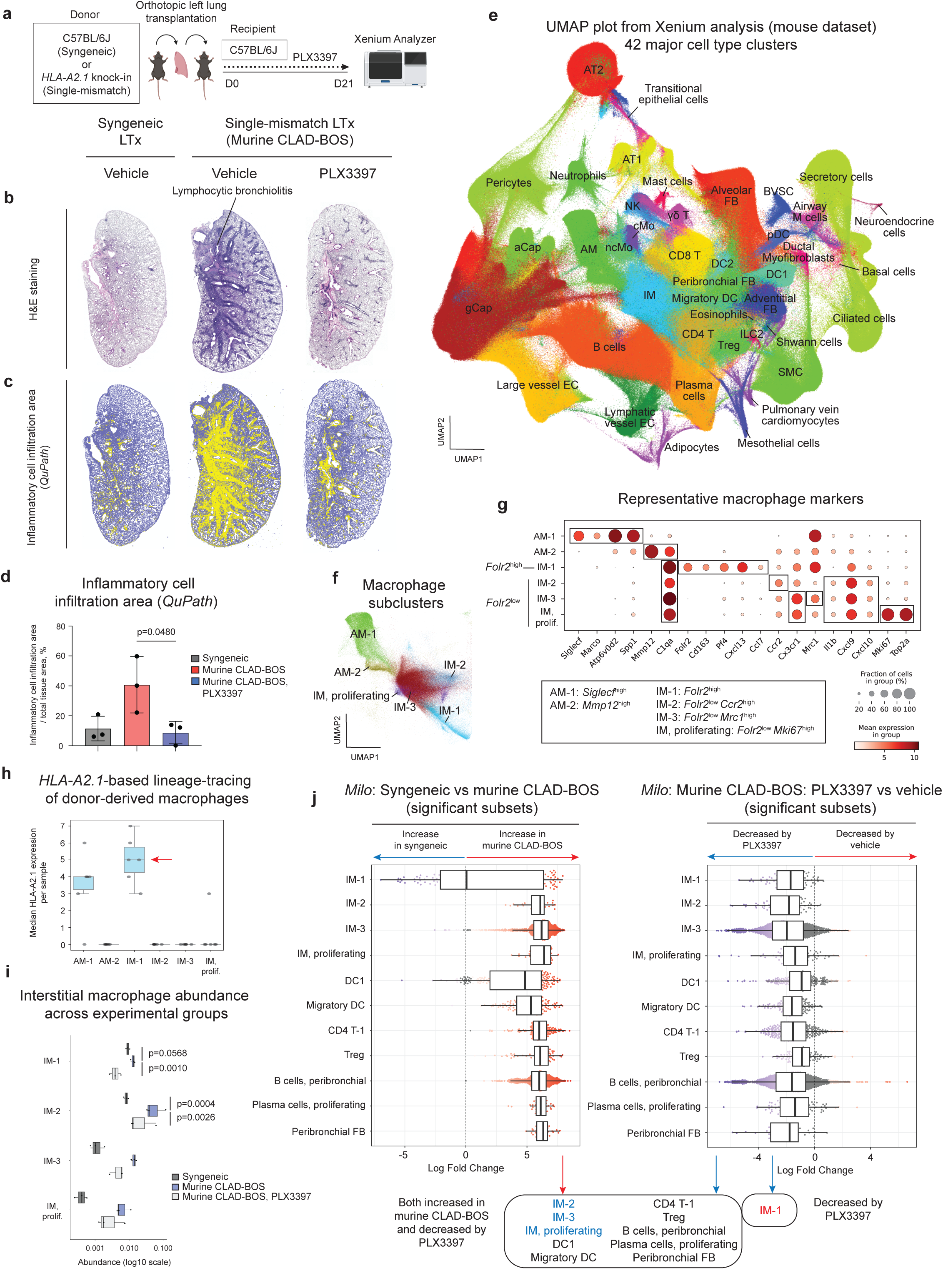
CSF1R blockade ameliorates CLAD-BOS pathology. **a.** Schematic of the experimental design for murine lung transplantation. Transplanted mice were treated daily with PLX3397 (40mg/Kg, orally) or vehicle and euthanized on day 21 for lung harvest. **b.** Representative hematoxylin and eosin (H&E)-stained lung allograft and syngeneic graft sections 21 days after transplantation. LTx = lung transplantation. **c.** Representative images highlighting the areas of inflammation identified using *QuPath* software. Yellow indicates positive regions (inflammatory cell infiltration) and blue indicates negative regions. **d.** Quantitative analysis of the proportion of inflammatory cell area relative to total tissue area identified using *QuPath* across experimental groups. Box plots represent the mean and standard deviation. Statistical significance was assessed using a one-way Analysis of Variance (ANOVA) followed by Tukey’s multiple comparisons test. ANOVA, p = 0.0381. **e.** UMAP visualization of single-cell spatial transcriptomic data from murine experiments (total number of cells = 3,502,604 cells, 479 genes, 42 major cell type clusters). Lung allografts and syngeneic grafts from three experimental groups (n = 3 per group; total n = 9) were included in this UMAP. aCap = alveolar capillary endothelial cells; AM = alveolar macrophages; AT1 = alveolar epithelial type I cells; AT2 = alveolar epithelial type II cells; BVSC = bronchovascular bundle sheath cells; CD4 T = CD4 T cells; CD8 T = CD8 T cells; cMo = classical monocytes; DC = dendritic cells; EC = endothelial cells; FB = fibroblasts; gCap = general capillary endothelial cells; γδ T = gamma delta T cells; ILC2 = group 2 innate lymphoid cells; IM = interstitial macrophages; ncMo = non-classical monocytes; NK = natural killer cells; pDC = plasmacytoid dendritic cells; SMC = smooth muscle cells; Treg = regulatory T cells. **f.** UMAP visualization of macrophage subsets. **g.** Dot plot showing expression of genes distinguishing each macrophage subtype. *Folr2*^high^ IM = IM-1; *Folr2*^low^ IM = IM-2, 3, and proliferating. **h.** *HLA-A2.1* expression across macrophage subtypes. Samples from single-mismatch lung transplantations (n = 6) were included in this analysis. Box plots represent the median and interquartile range. **i.** Differential abundance analysis of interstitial macrophage subsets based on per-sample fractions. Box plots represent the median and interquartile range. Statistical significance was assessed using pairwise *t*-tests with Bonferroni correction for multiple comparisons. **j.** Differential abundance analysis using *Milo* showing cell subsets significantly enriched in murine CLAD-BOS compared to syngeneic grafts with or without PLX3397 treatment. Box plots represent the median and interquartile range.

We used single-cell spatial transcriptomics to quantitatively characterize the response to CSF1R blockade in the mouse model of CLAD-BOS. We resolved 42 major cell types and 57 cell states, including rare cell populations absent in public scRNA-seq atlases,^4^ such as airway microfold cells (airway M cells), pulmonary vein cardiomyocytes, neuroendocrine cells, and others (**Figure 5e**, Suppl. Figure S9a-S9h). We identified two types of alveolar macrophages and four types of interstitial macrophages in mouse lungs (**Figure 5f**, 5g). Alveolar macrophage-1 (mouse AM-1), characterized by expression of *Siglecf*, matched homeostatic tissue-resident alveolar macrophages, whereas mouse AM-2 subset, characterized by *Mmp12* expression, corresponded to monocyte-derived alveolar macrophages.^25^ Within the interstitial macrophages, we resolved *Folr2*^high^ (mouse IM-1) and *Folr2*^low^ (mouse IM-2-4) subsets, mirroring observations in human data (**Figure 5f**, 5g). Mouse *Folr2*^low^ interstitial macrophages expressed *Cxcl9* and *Cxcl10*, similar to human *FOLR2*^low^ interstitial macrophages (**Figure 5g**). All mouse interstitial macrophage subsets were detected in every sample (Suppl. Figure S11a). Collectively, these data represent a comprehensive atlas of the mouse CLAD-BOS pathology and the therapeutic response to CSF1R-blockade.

### Lineage-tracing identifies origins of interstitial macrophage subsets in murine CLAD-BOS

We used expression of the *HLA-A2.1* transgene across all cell types as a genetic lineage-tag to distinguish cell ontogeny (**Suppl. Figure S10**). As expected, donor-derived structural cells, including epithelial, stromal, and endothelial cells, were *HLA-A2.1*-positive. In contrast, the majority of immune cells were *HLA-A2.1*-negative and thus recipient-derived. Notably, among the four interstitial macrophage subsets, only the *Folr2*^high^ interstitial macrophages (mouse IM-1) were *HLA-A2.1*-positive, confirming the donor-derived origin, whereas the remaining subsets were recipient monocyte-derived (**Figure 5h**).

Differential abundance analysis of per-sample cell fractions demonstrated an expansion of *Folr2*^high^ (mouse IM-1) and *Folr2*^low^ (mouse IM-2) interstitial macrophages in murine CLAD-BOS, both of which were significantly diminished following PLX3397 administration (**Figure 5i, Suppl. Figure S11b**). Next, we used *Milo* to further identify cell populations enriched in murine CLAD-BOS and significantly reduced upon PLX3397 administration (**Figure 5j**, **Suppl. Figure S12a, b**). The abundance of *Folr2*^high^ interstitial macrophage subset (mouse IM-1) was increased in CLAD-BOS and was significantly reduced by PLX3397. Similarly, *Folr2*^low^ interstitial macrophage subsets (mouse IM-2, 3, proliferating) were increased in CLAD-BOS and depleted by PLX3397. Notably, CSF1R-negative cell populations, including specific subsets of CD4 T cells, Tregs, B cells, and plasma cells, also increased in murine CLAD-BOS and decreased following PLX3397 treatment, suggesting that CSF1R-expressing interstitial macrophages organize lymphoid cell niches (**Figure 5j**). Among the fibroblast populations, only peribronchial fibroblasts increased in murine CLAD-BOS (**Suppl. Figure S12a**). Interestingly, this population was also reduced following PLX3397 administration (**Suppl. Figure S12b**). These results suggest that interstitial macrophages from both the donor lung and recipient are necessary to organize inflammatory infiltrates and expand activated fibroblast populations in a murine model of CLAD-BOS.

### *Folr2*^high^ interstitial macrophages are the predominant macrophage subset in perivascular adventitial and airway structural niches in murine syngeneic grafts

Next, we identified 9 multicellular spatial niches corresponding to microanatomical structures relevant to murine CLAD-BOS pathology. These spatial niches comprised the perivascular adventitial niche, perivascular immune niche, peribronchial TLS niche, airspace niche, airway epithelial niche, airway structural niche, alveolar niche, arterial niche, and distal arteriolar niche (**Figure 6a, Suppl. Figure S13a, b**). These niches, along with their cellular composition and organization, were similar between mice and humans. The perivascular adventitial niche was characterized by the accumulation of interstitial macrophages, adventitial fibroblasts, and mesothelial cells. The perivascular immune niche featured a dense aggregation of immune populations, including interstitial macrophage and DC subsets, alongside various lymphocyte subsets such as CD4 T cells, regulatory T cells (Treg), and group 2 innate lymphoid cells (ILC2). The peribronchial TLS niche shared a similar immune composition with the perivascular immune niche but was distinguished by a pronounced enrichment of B cells and plasma cells. The similarity between these two immune niches was confirmed by hierarchical clustering analysis of niche composition (**Figure 6b**). The airspace niche was dominated by alveolar macrophages and other immune cells. The airway epithelial, airway structural, and alveolar niches represented spatial structural units characterized by niche-specific combinations of epithelial, endothelial, stromal, and immune cells. While both the arterial and distal arteriolar niches were characterized by large-vessel endothelial cells, the distal arteriolar niche comprised distal vessels that lack smooth muscle cells (**Figure 6f**).

**Figure 6.**
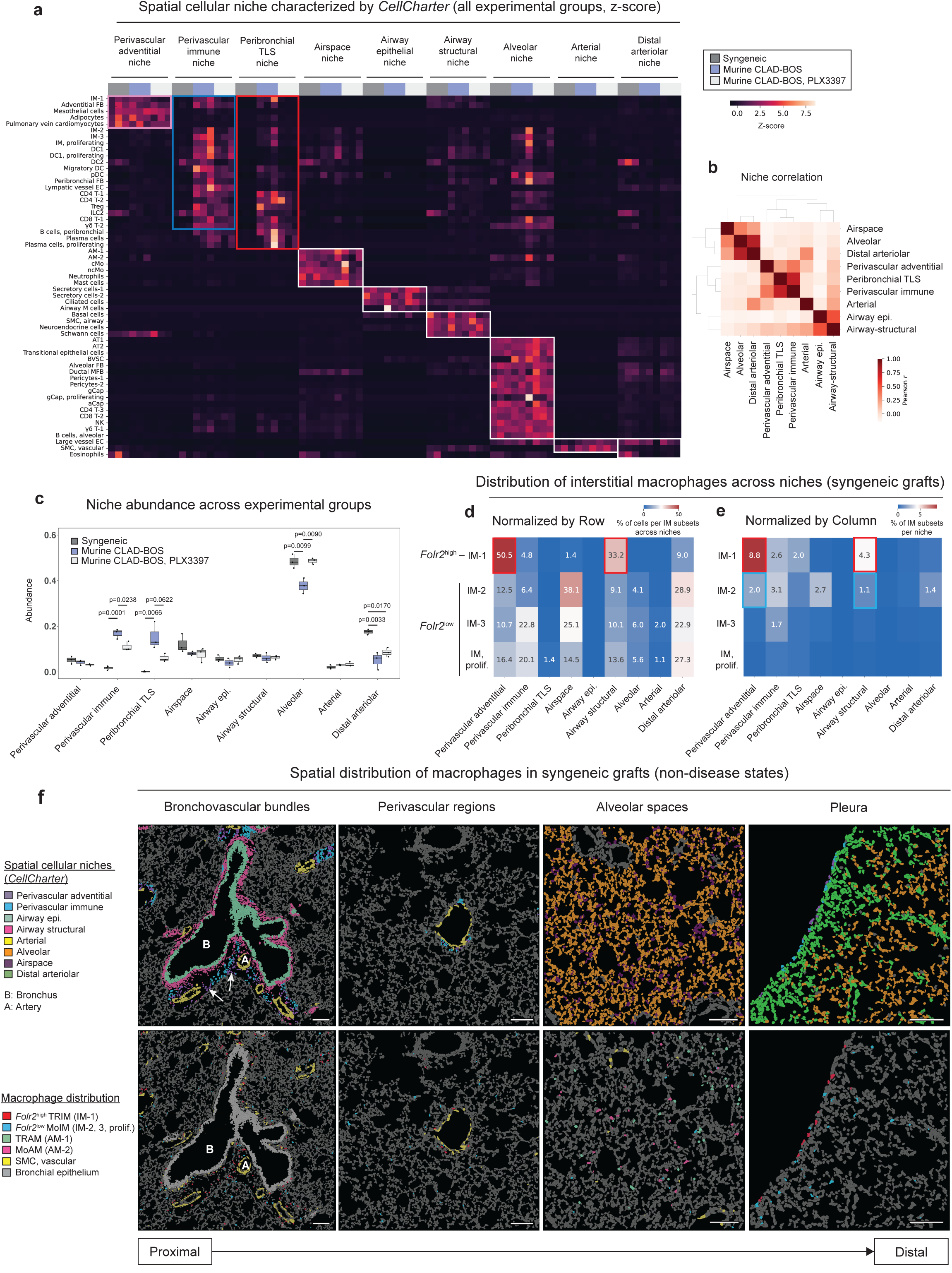
*Folr2*^high^ interstitial macrophages are the predominant macrophage subset in perivascular adventitial and airway structural niches in murine syngeneic grafts. **a.** Heatmap illustrating the enrichment of specific cell populations within each niche (z-score). **b.** Pearson correlation matrix showing similarity between niches based on their composition. **c.** Comparison of niche abundance across experimental groups. Box plots represent the median and interquartile range. Statistical significance was assessed using pairwise *t*-tests with Bonferroni correction for multiple comparisons. **d.** Distribution of interstitial macrophages across niches in syngeneic grafts (normalized by row). **e.** Distribution of interstitial macrophages across niches in syngeneic grafts (normalized by column). The heatmap displays the proportion of cells in each interstitial macrophage subset across niches, calculated by pooling cells from all samples. Values representing less than 1% are not shown. **f.** Representative images showing spatial cellular niches and the distribution of macrophage subsets across the proximal-to-distal axis of syngeneic grafts. Selected cell types are highlighted in each panel. TRIM = tissue-resident interstitial macrophages; MoIM = monocyte-derived interstitial macrophages; TRAM = tissue-resident alveolar macrophages, MoAM = monocyte-derived alveolar macrophages. Scale bars 100 μm.

In syngeneic grafts (non-disease states), the *Folr2*^high^ interstitial macrophage subset (mouse IM-1) represented the majority of interstitial macrophages within the perivascular adventitial and airway structural niches (perivascular adventitial niche: *Folr2*^high^ 78.8% vs *Folr2*^low^ 21.2%; airway structural niche: *Folr2*^high^ 76.4% vs *Folr2*^low^ 23.6%), whereas *Folr2*^low^ subsets (mouse IM-2, 3, proliferating) were sparsely distributed (**Figure 6d, 6e**). These spatial distributions were further confirmed via spatial mapping (**Figure 6f**). Together, these findings reveal that *Folr2*^high^ subset constitutes the majority of interstitial macrophages within bronchovascular bundles under non-diseased control conditions, consistent with observations in human donor lungs.

### CSF1R-expressing interstitial macrophages organize pathogenic niches in murine CLAD-BOS

The quantitative evaluation of niche abundance revealed that both the perivascular immune and peribronchial TLS niches were expanded in murine CLAD-BOS and reduced by PLX3397 treatment, suggesting that these pathogenic niches are regulated by CSF1R-expressing interstitial macrophages (**Figure 6c**). This is further supported by the significant alteration in the spatial distribution of interstitial macrophages observed in murine CLAD-BOS (**Figure 7a, 7b**). Specifically, in the context of murine CLAD-BOS, *Folr2*^high^ interstitial macrophages (mouse IM-1) were evenly distributed between the perivascular immune and peribronchial TLS niches, whereas *Folr2*^low^ interstitial macrophage subsets (mouse IM-2, 3, proliferating) were more frequently localized to the perivascular immune niche (**Figure 7a**). Interestingly, mirroring the detection of interstitial macrophages in human bronchoalveolar lavage fluid, we detected interstitial macrophages in the alveolar space.^18,19^ Recipient-derived *Folr2*^low^ interstitial macrophages were more frequent in the alveolar space than donor-derived *Folr2*^high^ interstitial macrophages (**Figure 7a, 7b**). PLX3397 treatment did not change the distribution of interstitial macrophage subsets, but it reduced their abundance within each niche (**Figure 7c-f**). The distribution of murine CLAD-BOS niches within bronchovascular bundles mirrored that observed in human CLAD-BOS lungs **(Suppl. Figure S14)**, and spatial mapping confirmed that PLX3397 treatment reduced both perivascular immune and peribronchial TLS niches and their constituent cells (**Figure 7g**).

**Figure 7.**
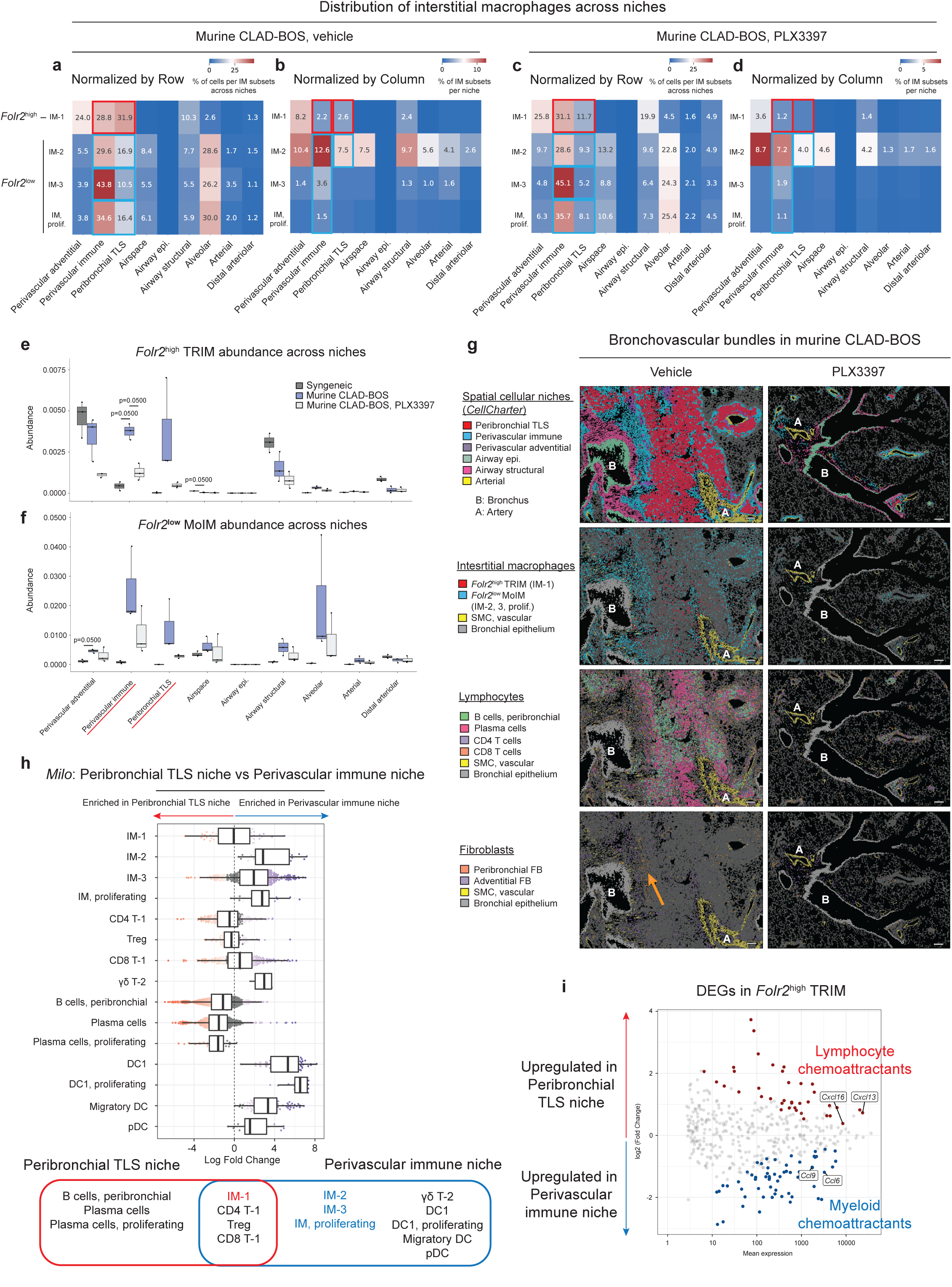
CSF1R-expressing interstitial macrophages organize pathogenic niches in murine CLAD-BOS. **a, c.** Distribution of interstitial macrophages across niches in murine CLAD-BOS lungs (**a**) and those treated with PLX3397 (**c**) (normalized by row). **b, d.** Distribution of interstitial macrophages across niches in murine CLAD-BOS lungs (**b**) and those treated with PLX3397 (**d**) (normalized by column). The heatmap displays the proportion of cells in each interstitial macrophage subset across niches, calculated by pooling cells from all samples. Values representing less than 1% are not shown. **e, f.** Relative abundance of *Folr2*^high^ (**e**) and *Folr2*^low^ (**f**) interstitial macrophages across niches and experimental groups. Box plots represent the median and interquartile range. **g.** Representative images highlighting the distribution of spatial cellular niches and their constituent cell types within the bronchovascular bundles of murine CLAD-BOS lungs and those treated with PLX3397. Arrow indicates the accumulation of peribronchial fibroblasts (FB) around the bronchi. Selected cell types are highlighted in each panel. Scale bars 100 μm. **h.** Differential abundance analysis using *Milo* showing significantly enriched immune cell subsets in the peribronchial TLS niche compared to the perivascular immune niche in murine CLAD-BOS lungs. Box plots represent the median and interquartile range. **i.** MA plot showing DEGs between *Folr2*^high^ interstitial macrophages in the peribronchial TLS niche and *Folr2*^high^ interstitial macrophages in the perivascular immune niche. DEGs with a |log_2_(fold-change)| > 0.585 (representing a ≥ 1.5-fold change in expression) and an adjusted *p*-value < 0.05 are plotted. Key representative markers are highlighted.

In addition to T and B cells, other cells identified in TLS structures were reduced in mice treated with PLX3397. Airway M cells are specialized mucosal epithelial cells that facilitate TLS formation by transporting antigens from the airway lumen to underlying immune cells, thereby initiating adaptive immune responses within bronchovascular bundles.^26,27^ In our murine CLAD-BOS analysis, we distinguished these cells, characterized by reported markers such as *Ccl9* and *Gp2*, from other epithelial cell populations and confirmed their spatial localization within the bronchial epithelium (**Suppl. Figure S9c, Figure S15**). In addition, we identified a distinct epithelial cell population localized at the interface between the bronchovascular bundle and the alveolar epithelium, which we designated as bronchovascular bundle sheath cells (BVSC) (**Suppl. Figure S15**). BVSC exhibited a transcriptional profile distinct from that of airway M cells, characterized by higher expression of alveolar epithelial-associated transcripts (*Ager* and *Lamp3*) and lower expression of bronchial epithelial markers (*Scgb3a2*, *Reg3g*, *Foxj1*, and others), while sharing certain signature genes, such as *Ccl9* and *Gpa33* (**Suppl. Figure S9c**). Among all epithelial cell populations analyzed, BVSC uniquely expressed *Aif1l* (**Suppl. Figure S9c**). Notably, both airway M cells and BVSC emerged in murine CLAD-BOS and were depleted following PLX3397 treatment (**Suppl. Figure S12a, S12b**), suggesting that their maintenance may rely on feedback from the cells localized within the TLS. Collectively, these results suggest that interstitial macrophages orchestrate the formation of multicellular TLS structures, including airway M cells and BVSC, in this murine model of CLAD-BOS.

### Donor-derived *Folr2*^high^ interstitial macrophages adopt niche-specific chemokine programs in murine CLAD-BOS

We took advantage of the observation that donor-derived *Folr2*^high^ interstitial macrophages occupy two pathologic niches to test whether their phenotype differs as a function of niche specific signals. We used *Milo* to show that while the distribution of donor-derived *Folr2*^high^ interstitial macrophages (mouse IM-1), CD4 and CD8 T cells, and Treg, was comparable across both niches, the peribronchial TLS niche was enriched for B cells and plasma cells (**Figure 7h**). In contrast, the perivascular immune niche was enriched for recipient-derived *Folr2*^low^ interstitial macrophages (mouse IM-2, 3, proliferating), γδ T cells, and DCs. Next, we performed differential gene expression analysis of donor-derived *Folr2*^high^ interstitial macrophages between these niches. Donor-derived *Folr2*^high^ tissue-resident interstitial macrophages in the peribronchial TLS niche expressed lymphocyte chemoattractants (*Cxcl13*, *Cxcl16*), while those in the perivascular immune niche predominantly expressed myeloid chemoattractants (*Ccl6*, *Ccl9*) (**Figure 7i**, **Suppl. Figure S16, Table S5**). This distinct expression pattern aligns with the cellular composition of each niche, suggesting that donor-derived tissue-resident interstitial macrophages adopt specialized functional states to orchestrate distinct immune microenvironments.

## DISCUSSION

Our study suggests that interstitial macrophages organize spatially restricted immune niches in the lungs of patients with CLAD-BOS. In a murine model of CLAD-BOS, pharmacological depletion of interstitial macrophages via CSF1R inhibition reduced TLS formation and fibroblast accumulation to attenuate allograft pathology. These findings implicate CSF1R-expressing interstitial macrophages in sustaining the macrophage– lymphocyte–fibroblast circuit associated with chronic allograft injury.^20,28^

We resolved conserved subsets of *FOLR2/Folr2* high- and low-interstitial macrophages in humans and mice.^21^ Notably, *FOLR2*/*Folr2*^high^ interstitial macrophages occupied perivascular stromal niches under homeostatic conditions in both species. In CLAD-BOS, these cells redistributed across expanded perivascular immune and peribronchial TLS niches and acquired niche-specific transcriptional programs. *Folr2*^high^ interstitial macrophages in peribronchial TLS niches preferentially expressed lymphocyte chemoattractants, whereas those in perivascular immune niches expressed chemokines associated with myeloid-cell recruitment. *FOLR2/Folr2*^low^ interstitial macrophages also expanded within CLAD-BOS bronchovascular bundles and expressed *CXCL9* and *CXCL10*, consistent with the recruitment or retention of CXCR3-expressing T cells. Thus, interstitial macrophage subsets exhibit spatially restricted cytokine programs reflecting the cellular composition of their local microenvironments.

Genetic lineage tracing combined with sensitive spatial techniques revealed distinct ontogenies for these interstitial macrophage subsets. In a murine model of CLAD-BOS, both donor-derived *Folr2*^high^ interstitial macrophages and recruited monocyte-derived *Folr2*^low^ interstitial macrophages contributed to the expansion of the interstitial macrophage pool. In contrast, in samples from patients with CLAD-BOS, most interstitial macrophages were recipient-derived, irrespective of *FOLR2* expression. This suggests that donor-derived interstitial macrophages are progressively replaced by recipient cells that acquire tissue-adapted phenotypes during the prolonged course of human disease. Alternatively, this difference may reflect species-specific macrophage kinetics, the compressed timescale of the murine model, or undersampling of interstitial macrophages during tissue dissociation. Systematic longitudinal sampling will be required to define the dynamics of macrophage replacement after transplantation.

Pharmacological targeting of CSF1R via PLX3397 implicates that interstitial macrophages actively support pathogenic, lymphocyte-rich niches in CLAD-BOS. Both donor-derived tissue-resident and recipient monocyte-derived interstitial macrophages were depleted by PLX3397, leading to the amelioration of lymphocytic bronchiolitis. Consistently, interstitial macrophages play an important role in mouse models of allergen- or viral lung injury-induced TLS formation^20^ and cGVHD.^16^ Since antibodies targeting CSF1R in humans are now available and have shown efficacy in patients with cGVHD,^13^ our findings provide support for clinical trials evaluating CSF1R blockade in patients with lung allograft rejection.

Our spatial atlas also identified BVSC, a previously unrecognized epithelial population at the interface between bronchovascular bundles and the alveolar parenchyma. BVSC expressed alveolar epithelial markers but shared selected transcriptional features with airway M cells, including *Ccl9* and *Gpa33*. Expression of some of these genes has been observed in intestinal M cells near Peyer’s patches.^29^ BVSC and airway M cells were enriched in allogeneic grafts and reduced after PLX3397 treatment, suggesting that their abundance is influenced by the inflammatory milieu. While we did not observe BVSC in our spatial analysis of samples from patients with CLAD-BOS, our targeted spatial transcriptomic panel was not designed to resolve them. Their ontogeny, function, and relationship to macrophage-dependent immune niches remain unresolved.

Several limitations should be considered. Human analyses were based predominantly on cross-sectional explant tissue reflecting advanced disease. Analysis of serial surveillance biopsies may close this gap and provide insights into the early origins of CLAD-BOS and its relationship to acute allograft rejection.^30^ Donor-recipient assignment was performed in a limited number of human samples, and targeted spatial panels constrained the unbiased analysis of transcriptional states and intercellular signaling. Although the affinity of PLX3397 for Fms-like tyrosine kinase 3 (FLT3) is nearly ten-fold lower than for CSF1R, there remains a potential for off-target effects on DCs, which were also reduced in PLX3397-treated animals.^31^ Finally, although spatial organization and chemokine expression are consistent with macrophage-mediated recruitment and retention of immune cells, these processes were not measured directly.

In summary, our findings identify macrophage-dependent immune remodeling as a driver of CLAD-BOS. Donor-derived tissue-resident and recipient monocyte-derived interstitial macrophages acquire complementary, niche-specific programs that support lymphoid and stromal accumulation. CSF1R inhibition disrupts these multicellular niches and attenuates allograft pathology, providing a rationale for evaluating macrophage-directed therapies in CLAD-BOS.

## METHODS

### Human subjects

All procedures involving human subjects were approved by the Northwestern University Institutional Review Board and the Office for Human Research Protections. Informed consent was obtained from all participants prior to tissue collection. Lung tissue samples were obtained from patients undergoing lung transplantation and from organ donors at Northwestern Medicine under study protocol STU00212120. The diagnoses of CLAD-BOS and IPF were clinically determined by through local multidisciplinary discussion based on current guidelines.^32,33^ The patient characteristics for this study are summarized in **Suppl. Table S1**. Lung tissues harvested at the time of transplantation were stored on ice until processing, fixed in 10% neutral-buffered formalin at 4°C for 24 hours, and then transferred to 70% ethanol. Formalin-fixed, paraffin-embedded (FFPE) tissue blocks were prepared using Sakura Tissue-Tek VIP Tissue Processor (Sakura Finetek USA, Inc.) at the Pathology Core Facility of the Robert H. Lurie Comprehensive Cancer Center at Northwestern University.

### Mice

All experimental protocols were approved by the Institutional Animal Care and Use Committee (IACUC) at Northwestern University (IS00015083). The C57BL/6J (B6, JAX:000664) and C57BL/6-*Mcph1^Tg(HLA–A2.1)1Enge^*/J (*HLA-A2.1* knock-in, JAX:003475) were obtained from the Jackson Laboratory. All strains were bred and housed at an Association for Assessment and Accreditation of Laboratory Animal Care International (AAALAC)-accredited facility under the administrative oversight of the Center for Experimental Animal Resources and the Animal Care and Use Committee. Veterinarian consultations are available, and a comprehensive training program ensures compliance with National Institutes of Health (NIH) Guidelines for the Care and Use of Laboratory Animals. The Northwestern University Center for Comparative Medicine operated a satellite facility adjacent to the laboratories for the temporary housing of experimental mice after their transfer from the main vivarium.

### Mouse lung transplantation, drug administration, and sample collection

Mice aged 8-14 weeks served as donors (C57BL/6J for syngeneic lung transplantation; *HLA-A2.1* knock-in for single-mismatch lung transplantation), while mice aged 10-17 weeks (C57BL/6J) were used as recipients. Orthotopic murine left lung transplantation was performed as previously described.^34–37^ Briefly, donor mice were anesthetized using a mixture of xylazine (10mg/Kg) and ketamine (100mg/Kg), and a thoracotomy was performed. The donor lungs were flushed with 3ml of sterile saline solution via the pulmonary artery. The heart-lung-block was excised and kept in a cooled preservative solution (4°C). The bronchus, pulmonary vein, and artery were dissected and prepared for anastomosis. A customized cuff made from a Teflon intravenous catheter was applied to the vascular structures and secured with a 10-0 nylon ligature. After placing a microvessel clip on the bronchus to prevent airway infiltration by the preservative solution, the graft was stored at 4°C for 90-120 minutes of cold ischemic time prior to implantation. Recipient mice received an intraperitoneal injection of xylazine (10mg/Kg), ketamine (100mg/Kg), and buprenorphine (0.1mg/kg) 30 minutes prior to the incision. Recipient mice were intubated and ventilated, and a left-sided thoracotomy was performed through the third intercostal space. The native lung was gently clamped and exteriorized from the thoracic cavity. The spaces between the artery, vein, and bronchus were dissected separately. The artery and vein were temporarily occluded using 8-0 nylon ligatures. Anastomoses were completed by securing each cuff with 10-0 nylon ligatures. The 8-0 ligatures were then released - first from the vein, followed by the artery – and the lung was inflated. The chest incision was closed, and the recipients were weaned from the ventilator once spontaneous respiration resumed. Postoperatively, recipient mice received a subcutaneous injection of sustained-release buprenorphine (1mg/Kg) and meloxicam (20mg/Kg). No immunosuppressive agents were administered postoperatively in any group.

PLX3397 (Pexidartinib) (#206178, MedKoo Biosciences, USA) was administered orally daily at a dose of 40mg/Kg, as previously described.^25^ Mice were euthanized on day 21 to harvest the lungs. Lung tissues harvested following perfusion were fixed in 10% neutral-buffered formalin at 4°C for 24 hours and then transferred to 70% ethanol. FFPE tissue blocks were prepared using Sakura Tissue-Tek VIP Tissue Processor (Sakura Finetek USA, Inc.) at the Mouse Histology and Phenotyping Laboratory of the Robert H. Lurie Comprehensive Cancer Center at Northwestern University.

### Single-cell spatial transcriptomics

#### Gene panel design

For the analysis of human samples, the Xenium Human Lung Gene Expression Panel (289 genes), pre-designed by 10x Genomics, along with a custom add-on panel (100 genes) was used (**Suppl. Table S2**). For the analysis of mouse samples, the Mouse Tissue Atlassing panel (379 genes), pre-designed by 10x Genomics, was used in combination with a custom add-on panel (100 genes, including *HLA-A2.1*) (**Suppl. Table S3**).

#### Sample preparation and data generation

For human sample analysis, lung tissues were obtained from patients with CLAD-BOS (n = 3), IPF (n=4), and donors (n = 4). For murine studies, lung tissues were collected from three experimental groups: syngeneic lung transplantation (n = 3), single-mismatch lung transplantation (murine CLAD-BOS, n = 3), and single-mismatch lung transplantation treated with PLX3397 (n = 3). FFPE tissue blocks were rehydrated in an ice bath for 30 min and sectioned at 5µm thickness using a HistoCore BIOCUT microtome (Leica Biosystems, USA). Mounted sections on Xenium slides were baked at 42 °C for 3 hours, followed by deparaffinization, de-crosslinking, probe hybridization, ligation, and amplification, according to the Xenium In Situ Gene Expression workflow (CG000754 Rev A). Xenium In Situ Cell Segmentation staining was performed using the Xenium Multi-Tissue Stain Mix (PN2000991), which provides four types of labeling: antibody-labeled membranes, antibody-labeled cell interiors, a universal interior label targeting ribosomal RNA, as well as the nuclear label 4’, 6-diamidino-2-phenylindole (DAPI). The data was generated on Xenium Analyzer (10x Genomics) using manufacturer protocol. Image data from the Xenium Analyzer were processed using the Xenium onboard analysis pipeline (version 3.2.0.7 for human samples and 3.0.0.15 for mouse samples).

#### Data processing

Off-instrument reanalysis of cell segmentation was performed using the Xenium Ranger (version 2.0.0.10, 10x Genomics). For each sample, Xenium generated an output file containing transcript information, including the x and y coordinates, corresponding gene target, assigned cell, a binary flag indicating whether the transcript was expressed over a nucleus, and a quality score. Transcript nucleus filtering was performed to retain high-quality signals (QV ≥ 20), and nuclear boundaries were expanded by 2 µm (or until hitting an adjacent boundary) using a Voronoi-based algorithm to define cellular limits. Seurat v5 was used to further quality filtering and visualization.^38^ A single merged Seurat object was created from all samples using region of interest (ROI) count matrices and metadata files containing nuclei coordinates and area information. Nuclei were retained according to the following criteria: ≥ 2 unique genes. Seurat v5 was then used to perform dimensionality reduction, clustering, and visualization. Gene expression was normalized per cell using Seurat’s *SCTransform* function.^39^ Dimensionality reduction was performed using principal component analysis (PCA) on all target genes. Cells were clustered using the Louvain algorithm based on the first 40 principal components (PCs). These PCs were utilized as input for Uniform Manifold Approximation and Projection (UMAP) to visualize the data in two dimensions. Cell clusters were then manually annotated using a combination of canonical marker gene expression and spatial context, including cell morphology and anatomical positioning within the tissue.

#### Spatial differential abundance analysis

Spatial differential abundance analyses were performed using two complementary approaches: 1) Per-sample fractions, in which cell abundance was analyzed by summing cells of each type within individual samples. These per-sample fractions were then compared across cell types using pairwise *t*-tests with Bonferroni correction for multiple comparisons. 2) *Milo*, a neighborhood-based differential abundance testing framework that models local changes in cell composition across experimental conditions.^22^ Briefly, *Milo* constructs a κ-nearest neighbor (κ-NN) graph of cells based on transcriptomics similarity and tests for differential abundance within graph-defined neighborhoods, enabling robust detection of subtle spatial or compositional shifts. To assess differential abundance, *Milo* applied a negative binomial generalized linear model (GLM) to neighborhood-level counts, using Trimmed Mean of M-values normalization to correct for variations in total cell numbers and compositional biases across samples. Cell populations were defined as significantly changed if at least four neighborhoods were identified, and both the upper and lower quartiles of log-fold change were skewed to one side relative to zero.

#### Spatial niche detection

Spatial niche analysis was performed using *CellCharter*, a computational framework that identifies cellular niches based on both transcriptomic similarity and spatial proximity.^23^ *CellCharter* integrates spatial coordinates with gene expression data to cluster cells into distinct niches, facilitating the characterization of local microenvironments and their compositional relationships. This approach enables the identification of spatially co-localized cell populations and the investigation of how niche-specific interactions differ across conditions. Niche abundance was analyzed by counting the number of cells assigned to each *CellCharter*-defined niche per sample, normalizing these counts to fractions to account for differences in total cell numbers, and comparing niche proportions across experimental conditions using pairwise *t*-tests with Bonferroni correction for multiple comparisons.

#### Spatial niche proximity analysis

To evaluate the spatial relationships between distinct niches, cellular coordinates (x and y positions corresponding to the centers of segmented cells) extracted from the Xenium Ranger outputs were used to define niche boundaries. The absolute distance from edge cells of each niche to the nearest cells of other distinct niches was then calculated for each sample.

#### Differential gene expression analysis

Spatial transcriptomics data generated by Xenium were analyzed for differential gene expression using *DESeq2*.^40^ Prior to differential expression analysis, data quality was assessed, and appropriate dispersion estimates were confirmed for the count data. Low-count genes were filtered to remove sparsely expressed targets, and counts were aggregated by cell type to reduce sparsity and account for the structure of spatially resolved data. These dispersion estimates indicate that *DESeq2* is suitable for detecting differentially expressed genes in our dataset. Differential expression between experimental conditions was performed using *DESeq2*’s GLM framework, which models count data with a negative binomial distribution. Resulting log2-fold changes and adjusted p-values (Benjamini-Hochberg correction) were used to identify significantly differentially expressed genes. To further filter out noise from segmentation artifacts, significant genes that were expressed in < 25% of cells in a given condition were removed from analysis.

### Integrated scRNA-seq analysis across independent, publicly available datasets

#### Integration

To complement our single-cell spatial analysis findings, we integrated and reanalyzed three independent, publicly available single-cell RNA-seq datasets of CLAD-BOS; Khatri *et al.* (GSE224210), Mellors *et al*. (GSE290834), and Yan *et al.* (GSE289881).^5,7,11^ Data integration and clustering were performed using single-cell variational inference (scVI) and *Scanpy* (version 1.9.8) as previously described.^19,41,42^ Low-quality cells were removed, and cell types were manually annotated for each cluster based on established marker genes.

#### Differential gene expression analysis

For differential gene expression, raw count matrices were aggregated into pseudobulk profiles by summing counts across cells of the same cell type within each sample. These pseudobulk count matrices were analyzed using *DESeq2*, which applies a negative binomial GLM with internal normalization for sequencing depth.^40^ Resulting log2-fold changes and adjusted p-values (Benjamini-Hochberg correction) were used to identify significantly differentially expressed genes. To interpret differential expression results, gene set enrichment analysis (GSEA) was performed on ranked genes for each cell type using DESeq2-estimated log_2_FoldChange. We utilized 50 hallmark gene sets from MsigDB, with FDR correction applied across all gene sets and cell types. The enrichment scores were visualized as -log_10_(*padj*) X sign (*ES*).

#### Genetic demultiplexing and donor-recipient assignment

To determine the genetic ontogeny of cells within our own dataset (Yan *et al.*; GSE289881), we performed genetic demultiplexing using *Souporcell*.^24^ Raw sequencing data (FASTQ files) were processed and aligned to the human reference genome (GRCh38) to generate binary alignment map (BAM) files. These were used as input for *Souporcell*, which identifies individual genotypes by clustering cells based on single-nucleotide polymorphisms (SNPs) found within the RNA-seq reads. By performing *de novo* variant calling, the algorithm distinguished cells into two distinct genotypes (κ = 2), representing the donor and recipient populations, without requiring reference genotypes. Cells identified as inter-genotypic doublets or those with unassignable genotypes were excluded from downstream analysis. The resulting clusters were assigned as ‘donor” or “recipient” based on the assumption that structural cells (e.g. epithelial cells and endothelial cells) are donor-derived in the context of lung transplantation.

#### Histopathological analysis of mouse lung sections

Mouse lung sections stained with Hematoxylin and Eosin were scanned as NDPI files (Hamamatsu Photonics K.K., Japan) and analyzed using *QuPath* (version 0.6.0, Queen’s University Belfast, UK).^43^ A custom image analysis algorithm was developed in *QuPath* to train and classify regions as Positive (inflammatory cell infiltration), Negative, or Blank. This algorithm was applied consistently across all samples. The proportion of inflammatory cell infiltration area relative to total tissue area was calculated for each sample.

### Statistical analysis

Sample size was determined based on sample availability and the number of samples that could be accommodated on a Xenium slide. Given the nature of the study, randomization and blinding were not performed. All statistical analyses are reported as described in the Method sections or Figure legends. A significance threshold of p < 0.05 was applied for all tests. Statistical analyses were performed using R (version 4.2.3), Python (version 3.8.19), or Prism (version 10, GraphPad, USA).

## Data availability

Raw and processed data are deposited at GEO under accession GSE343736 (mouse) and GSE343735 (human).

## Code availability

All code used for the analysis is available at https://github.com/NUPulmonary/2026_Suzuki_CLAD.

Spatial transcriptomics data can be explored via the data browsers at https://sqlifts.fsm.northwestern.edu/public/2026_Suzuki_CLAD/.

## Acknowledgments

We thank all patients who provided samples and data for this study. This research was supported in part by a generous gift from Kimberly Querrey and Louis A. Simpson, as well as by the Simpson Querrey Lung Institute for Translational Science (SQLIFTS) at Northwestern University. This work was supported by the Northwestern University Pathology Core Facility and a Cancer Center Support Grant (NCI CA060553). Comparative histopathology and molecular phenotyping services were provided by the Mouse Histology and Phenotyping Laboratory (MHPL, RRIS:SCR_017870) at Northwestern University, which is supported by National Cancer Institute (NCI) P30-CA060553 grant awarded to the Robert H. Lurie Comprehensive Cancer Center. This research was also supported in part by the computational resources and staff contributions of the Quest high-performance computing facility at Northwestern University, jointly supported by the Office of the Provost, the Office for Research, and Northwestern University Information Technology. Additional support was provided through the computational resources and staff contributions of the Genomics Compute Cluster, which is jointly supported by the Feinberg School of Medicine, the Center for Genetic Medicine, Feinberg’s Department of Biochemistry and Molecular Genetics, the Office of the Provost, the Office for Research, and Northwestern Information Technology. The Genomics Compute Cluster is part of Quest, Northwestern University’s high-performance computing facility, aimed at advancing research in genomics. Integrative genomic services were provided by the Metabolomics Core Facility at the Robert H. Lurie Comprehensive Cancer Center of Northwestern University.

A.S. was supported by the Cell Science Research Foundation, the Cugell Fellowship, and the Pulmonary Fibrosis Foundation Scholars Award. A.B. was supported by the National Institutes of Health (NIH) (grants P01HL169188, R01HL147290, R01HL145478, and R01HL147575). A.V.M. was supported by the NIH (grants U19AI135964, U19AI181102, P01AG049665, P01HL154998, P01HL169188, R01HL153312, R01HL158139, and R01ES034350), and research grants from AbbVie, Merck, and Incyte. G.R.S.B. was supported by the NIH (grants U19AI135964, U19AI181102, U54AG079754, P01AG049665, P01HL071643, R01HL147575, R01HL145478, R01HL147290, R01HL173940, and R01HL154686), the US Department of Veterans Affairs (I01CX001777), and the Simpson Querrey Lung Institute for Translational Sciences. The funders had no role in the study design, data collection and analysis, the decision to publish, or manuscript preparation.

