## Supplementary Information for "Interstitial macrophages drive chronic lung allograft dysfunction"

**Supplementary Figures**

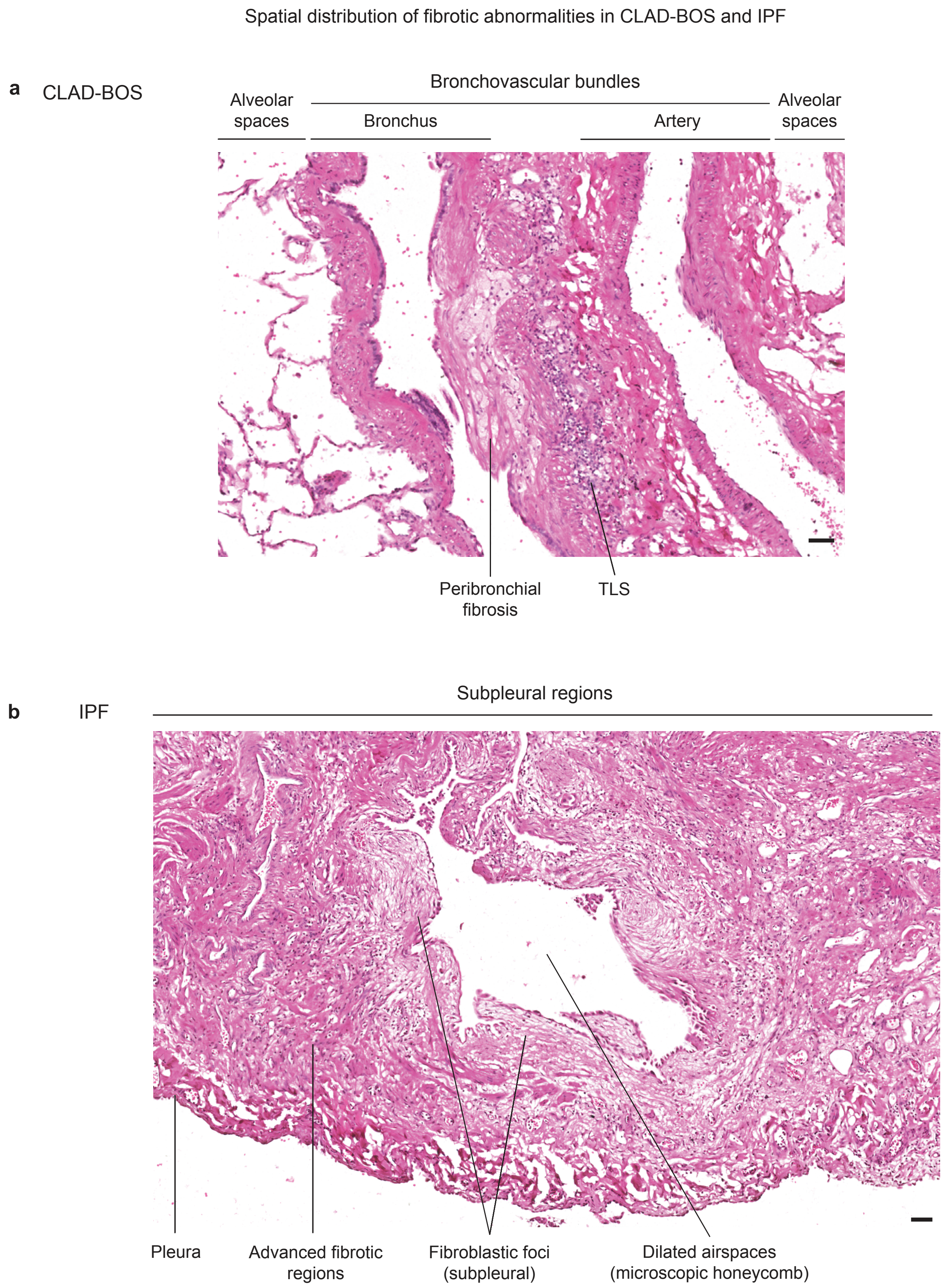


**Supplementary Figure S1. Spatial distribution of fibrotic abnormalities in CLAD-BOS and IPF lungs. a.** Representative hematoxylin and eosin-stained section from CLAD-BOS lung. This condition is characterized by inflammation and fibrosis around small airwaysin the bronchovascular bundle. TLS = tertiary lymphoid structures. **b.** Representative hematoxylin and eosin-stained section from IPF lung. This condition is characterized by progressive fibrosis in the subpleural alveolar space with relative sparing of the airways, and the presence of fibroblastic foci. Scale bars 50 μm.

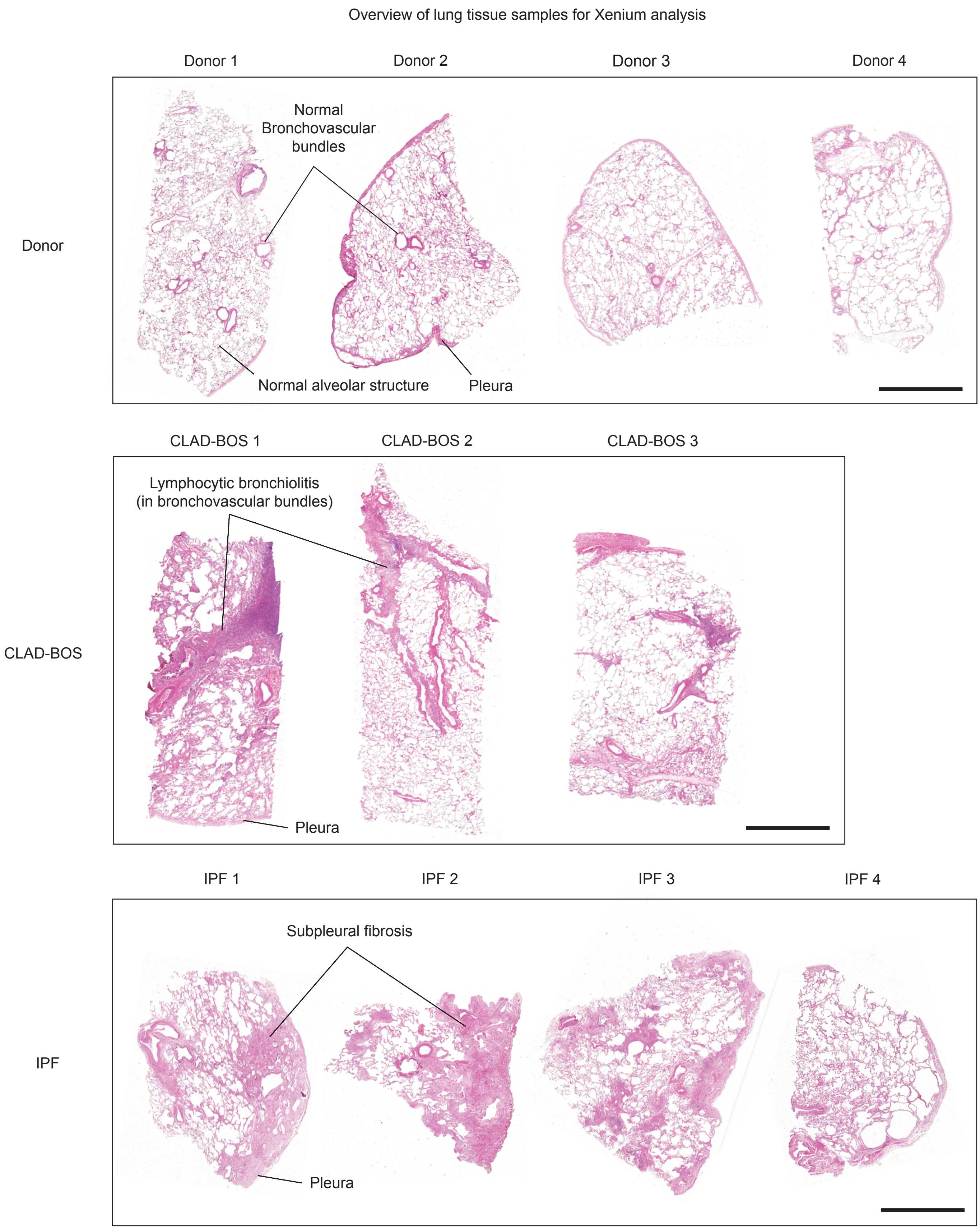


**Supplementary Figure S2. Hematoxylin and eosin staining of lung tissue sections used for single-cell spatial transcriptomic analysis.** Lung tissue from patients with CLAD-BOS (n = 3) and IPF (n = 4), and surgical biopsies from donor lungs accepted for transplantation (n = 4). Scale bars 2.5 mm.

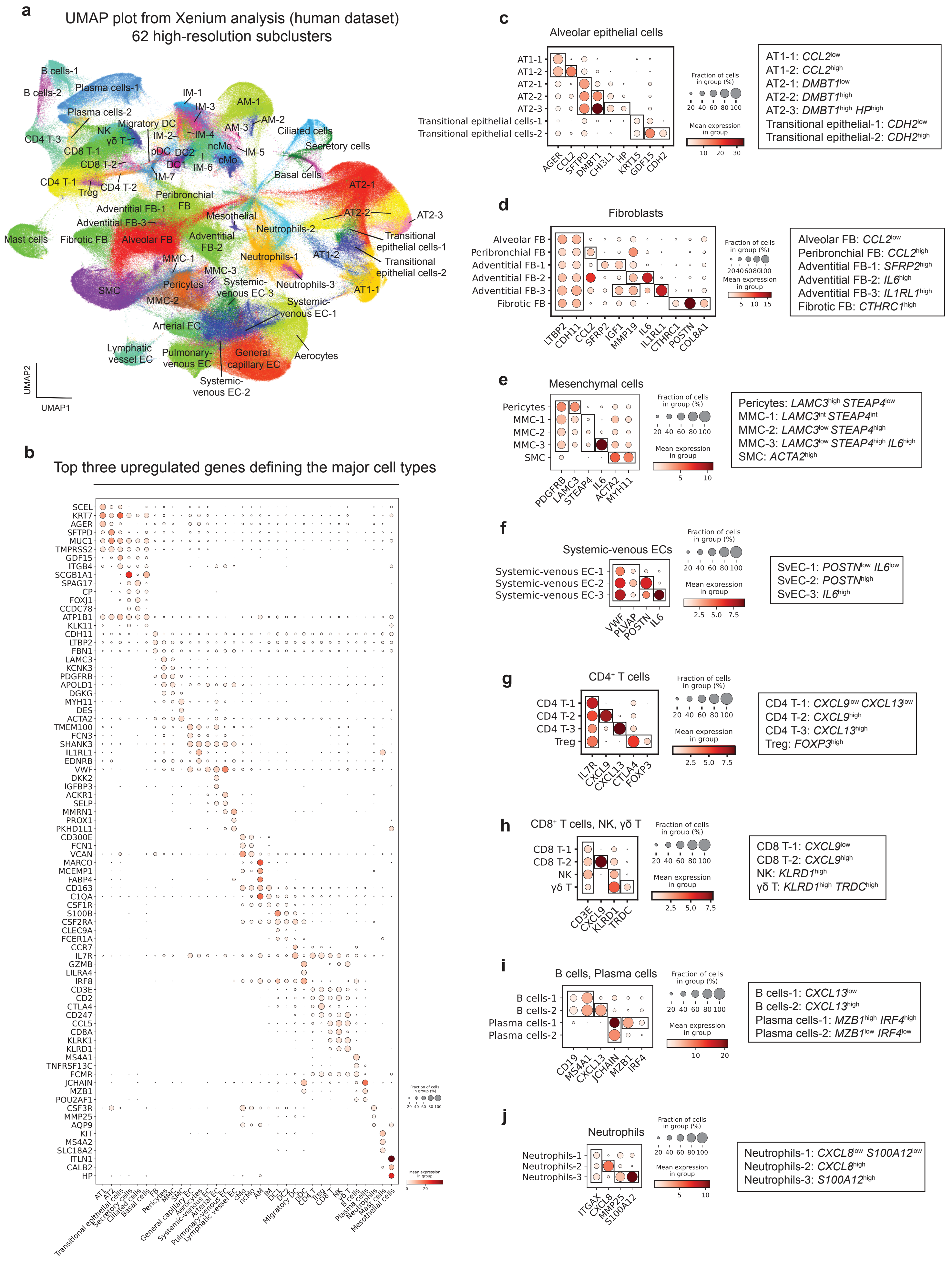

a
UMAP plot from Xenium analysis (human dataset)
62 high-resolution subclusters
B cells-1
B cells-2
Plasma cells-1
Plasma cells-2
IM-1
IM-3
AM-1
AM-2
AM-3
IM-2
IM-4
Migratory DC
NK
γδ T
CD4 T-3
CD8 T-1
CD8 T-2
pDC
DC2
DC1
IM-6
ncMo
cMo
IM-5
IM-7
Ciliated cells
Secretory cells
Basal cells
CD4 T-1
Treg
CD4 T-2
Peribronchial FB
Adventitial FB-1
Adventitial FB-3
Mesothelial
AT2-1
AT2-2
AT2-3
Mast cells
Fibrotic FB
Alveolar FB
Adventitial FB-2
Neutrophils-2
Transitional epithelial cells-1
Transitional epithelial cells-2
MMC-1
Neutrophils-1
AT1-2
MMC-3
Pericytes
Systemic-venous EC-3
Neutrophils-3
AT1-1
SMC
MMC-2
Systemic-venous EC-1
Arterial EC
Lymphatic vessel EC
Pulmonary-venous EC
General capillary EC
Aerocytes
Systemic-venous EC-2
UMAP2
UMAP1
b
Top three upregulated genes defining the major cell types
SCEL
KRT7
AGER
SFTPD
MUC1
TMPRSS2
GDF15
ITGB4
SCGB1A1
SPAG17
CP
FOXJ1
CCDC78
ATP1B1
KLK11
CDH11
LTBP2
FBN1
LAMC3
KCNK3
PDGFRB
APOLD1
DGKG
MYH11
DES
ACTA2
TMEM100
FCN3
SHANK3
IL1RL1
EDNRB
VWF
DKK2
IGFBP3
ACKR1
SELP
MMRN1
PROX1
PKHD1L1
CD300E
FCN1
VCAN
MARCO
MCEMP1
FABP4
CD163
C1QA
CSF1R
S100B
CSF2RA
CLEC9A
FCER1A
CCR7
IL7R
GZMB
LILRA4
IRF8
CD3E
CD2
CTLA4
CD247
CCL5
CD8A
KLRK1
KLRD1
MS4A1
TNFRSF13C
FCMR
JCHAIN
MZB1
POU2AF1
CSF3R
MMP25
AQP9
KIT
MS4A2
SLC18A2
ITLN1
CALB2
HP
AT1
AT2
Transitional epithelial cells
Secretory cells
Ciliated cells
Basal cells
FB
Pericytes
MMC
SMC
General capillary EC
Aerocytes
Systemic-venous EC
Arterial EC
Pulmonary-venous EC
Lymphatic vessel EC
cMo
ncMo
AM
IM
DC1
DC2
Migratory DC
pDC
CD4 T
Treg
CD8 T
NK
γδ T
B cells
Plasma cells
Neutrophils
Mast cells
Mesothelial cells
Fraction of cells in group (%)
20 40 60 80 100
Mean expression in group
0 20
c
Alveolar epithelial cells
AT1-1
AT1-2
AT2-1
AT2-2
AT2-3
Transitional epithelial cells-1
Transitional epithelial cells-2
AGER
CCL2
SFTPD
DMBT1
CHI3L1
HP
KRT15
GDF15
CDH2
Fraction of cells in group (%)
20 40 60 80 100
Mean expression in group
10 20 30
AT1-1: CCL2low
AT1-2: CCL2high
AT2-1: DMBT1low
AT2-2: DMBT1high
AT2-3: DMBT1high HPhigh
Transitional epithelial-1: CDH2low
Transitional epithelial-2: CDH2high
d
Fibroblasts
Alveolar FB
Peribronchial FB
Adventitial FB-1
Adventitial FB-2
Adventitial FB-3
Fibrotic FB
LTBP2
CDH11
CCL2
SFRP2
IGF1
MMP19
IL6
IL1RL1
CTHRC1
POSTN
COL8A1
Fraction of cells in group (%)
20406080100
Mean expression in group
5 10 15
Alveolar FB: CCL2low
Peribronchial FB: CCL2high
Adventitial FB-1: SFRP2high
Adventitial FB-2: IL6high
Adventitial FB-3: IL1RL1high
Fibrotic FB: CTHRC1high
e
Mesenchymal cells
Pericytes
MMC-1
MMC-2
MMC-3
SMC
PDGFRB
LAMC3
STEAP4
IL6
ACTA2
MYH11
Fraction of cells in group (%)
20 40 60 80 100
Mean expression in group
5 10
Pericytes: LAMC3high STEAP4low
MMC-1: LAMC3int STEAP4int
MMC-2: LAMC3low STEAP4high
MMC-3: LAMC3low STEAP4high IL6high
SMC: ACTA2high
f
Systemic-venous ECs
Systemic-venous EC-1
Systemic-venous EC-2
Systemic-venous EC-3
VWF
PLVAP
POSTN
IL6
Fraction of cells in group (%)
20 40 60 80 100
Mean expression in group
2.5 5.0 7.5
SvEC-1: POSTNlow IL6low
SvEC-2: POSTNhigh
SvEC-3: IL6high
g
CD4+ T cells
CD4 T-1
CD4 T-2
CD4 T-3
Treg
IL7R
CXCL9
CXCL13
CTLA4
FOXP3
Fraction of cells in group (%)
20 40 60 80 100
Mean expression in group
2.5 5.0 7.5
CD4 T-1: CXCL9low CXCL13low
CD4 T-2: CXCL9high
CD4 T-3: CXCL13high
Treg: FOXP3high
h
CD8+ T cells, NK, γδ T
CD8 T-1
CD8 T-2
NK
γδ T
CD3E
CXCL9
KLRD1
TRDC
Fraction of cells in group (%)
20 40 60 80 100
Mean expression in group
2.5 5.0 7.5
CD8 T-1: CXCL9low
CD8 T-2: CXCL9high
NK: KLRD1high
γδ T: KLRD1high TRDChigh
i
B cells, Plasma cells
B cells-1
B cells-2
Plasma cells-1
Plasma cells-2
CD19
MS4A1
CXCL13
JCHAIN
MZB1
IRF4
Fraction of cells in group (%)
20 40 60 80 100
Mean expression in group
10 20
B cells-1: CXCL13low
B cells-2: CXCL13high
Plasma cells-1: MZB1high IRF4high
Plasma cells-2: MZB1low IRF4low
j
Neutrophils
Neutrophils-1
Neutrophils-2
Neutrophils-3
ITGAX
CXCL8
MMP25
S100A12
Fraction of cells in group (%)
20 40 60 80 100
Mean expression in group
5 10
Neutrophils-1: CXCL8low S100A12low
Neutrophils-2: CXCL8high
Neutrophils-3: S100A12high

**Supplementary Figure S3. Single-cell spatial transcriptomics resolves cell types in the human lung. a.** Uniform manifold approximation and projection (UMAP) visualization of single-cell spatial transcriptomic data from human samples (total number of cells = 881,761 cells, 389 genes, 62 high-resolution subclusters). AM = alveolar macrophages; AT1 = alveolar epithelial type I cells; AT2 = alveolar epithelial type II cells; CD4 T = CD4 T cells; CD8 T = CD8 T cells; cMo = classical monocytes; DC = dendritic cells; EC = endothelial cells; FB = fibroblasts; γδ T = gamma delta T cells; IM = interstitial macrophages; MMC = microvascular mural cells; ncMo = non-classical monocytes; NK = natural killer cells; pDC = plasmacytoid dendritic cells; SMC = smooth muscle cells; Treg = regulatory T cells. **b.** Dot plot showing the top three marker genes for each major cell type. **c.** Dot plot showing selected genes used to resolve alveolar epithelial cell types. **d.** Dot plot showing selected genes used to resolve fibroblast subtypes. **e.** Dot plot showing selected genes used to resolve mesenchymal subsets. **f.** Dot plot showing selected genes used to resolve systemic-venous endothelial cells (SvEC). **g.** Dot plot showing selected genes used to resolve CD4 T cells and Treg. **h.** Dot plot showing selected genes used to resolve CD8 T cells, NK cells, and γδ T cells. **i.** Dot plot showing selected genes used to resolve B cells and plasma cells. **j.** Dot plot showing selected genes used to resolve neutrophils.

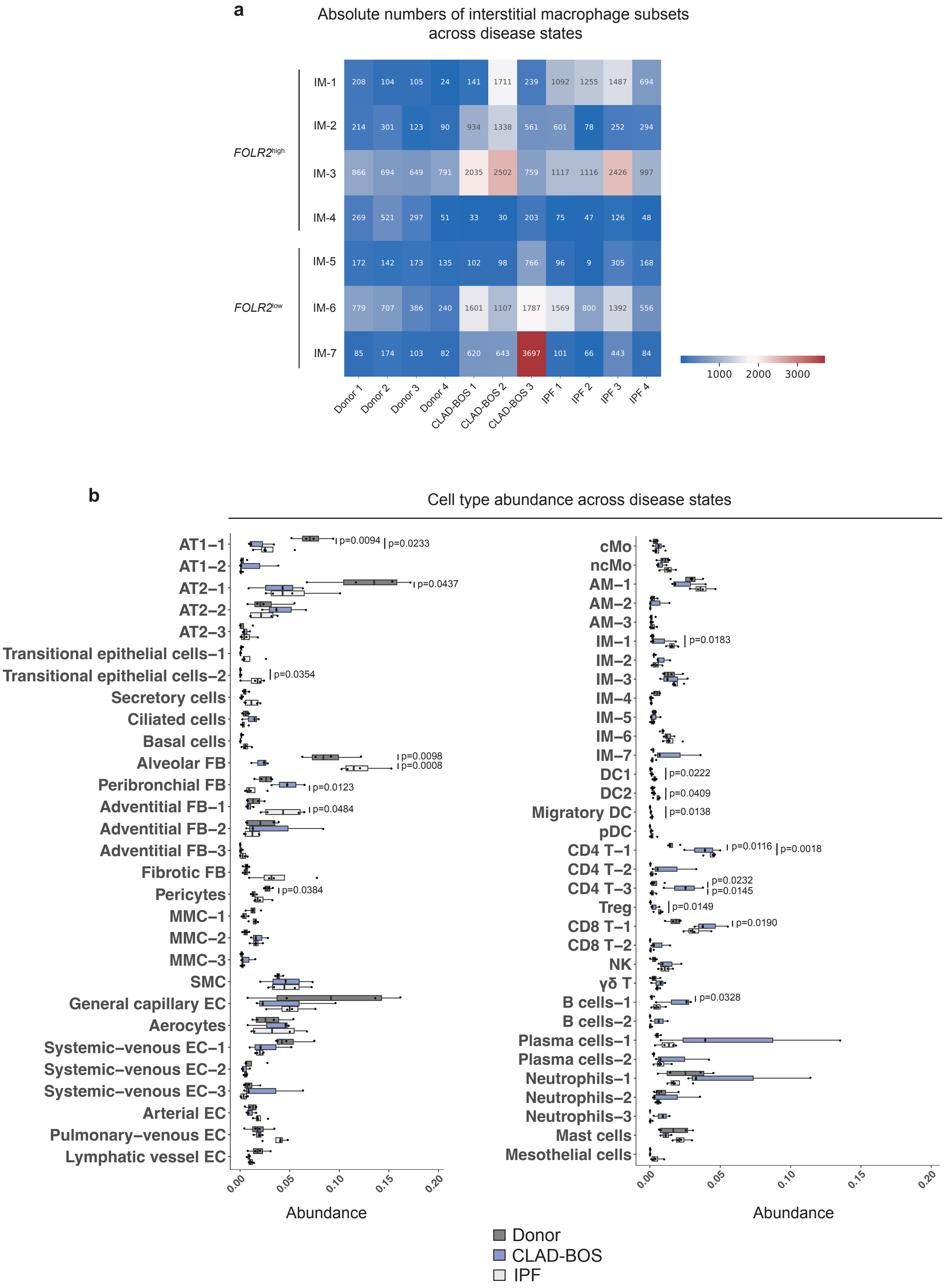


**Supplementary Figure S4. Cell type abundance between donor, CLAD-BOS, and IPF lungs using per-sample fractions. a.** Absolute numbers of interstitial macrophages across disease states. *FOLR2*$^{high}$ IM = IM-1-4; *FOLR2*$^{low}$ IM = IM-5-7. **b.** Differential abundance analysis based on per-sample fractions. Box plots represent the median and interquartile range. Statistical significance was assessed using pairwise t-tests with Bonferroni correction for multiple comparisons.

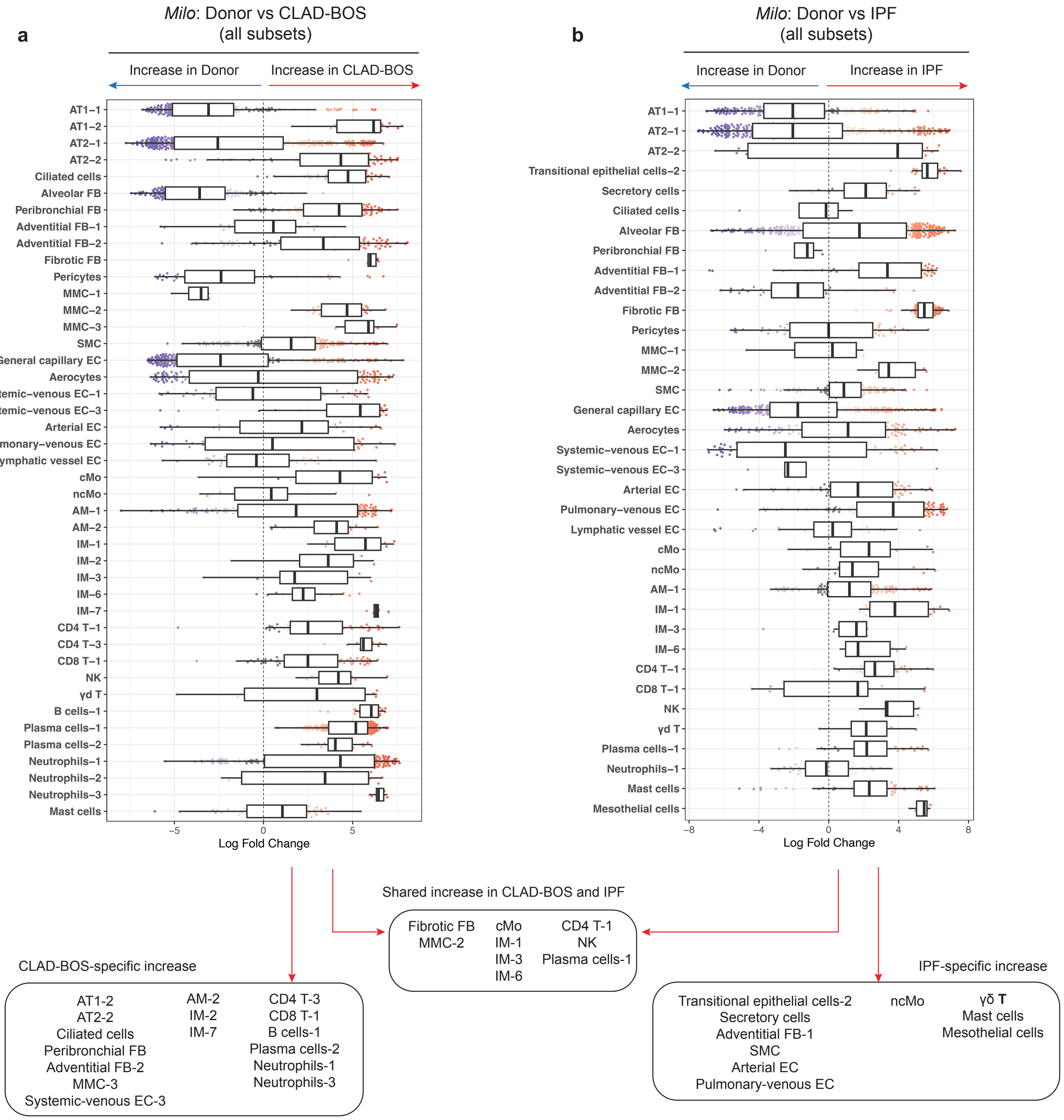


**Supplementary Figure S5. Cell type abundance between donor, CLAD-BOS, and IPF lungs using *Milo*. a, b.** Differential abundance analysis using *Milo* showing all cell types in CLAD-BOS and IPF compared to donor lungs. Box plots represent the median and interquartile range. Cell populations were defined as significantly changed if at least four neighborhoods were identified, and both the upper and lower quartiles of log-fold change were skewed to one side relative to zero.

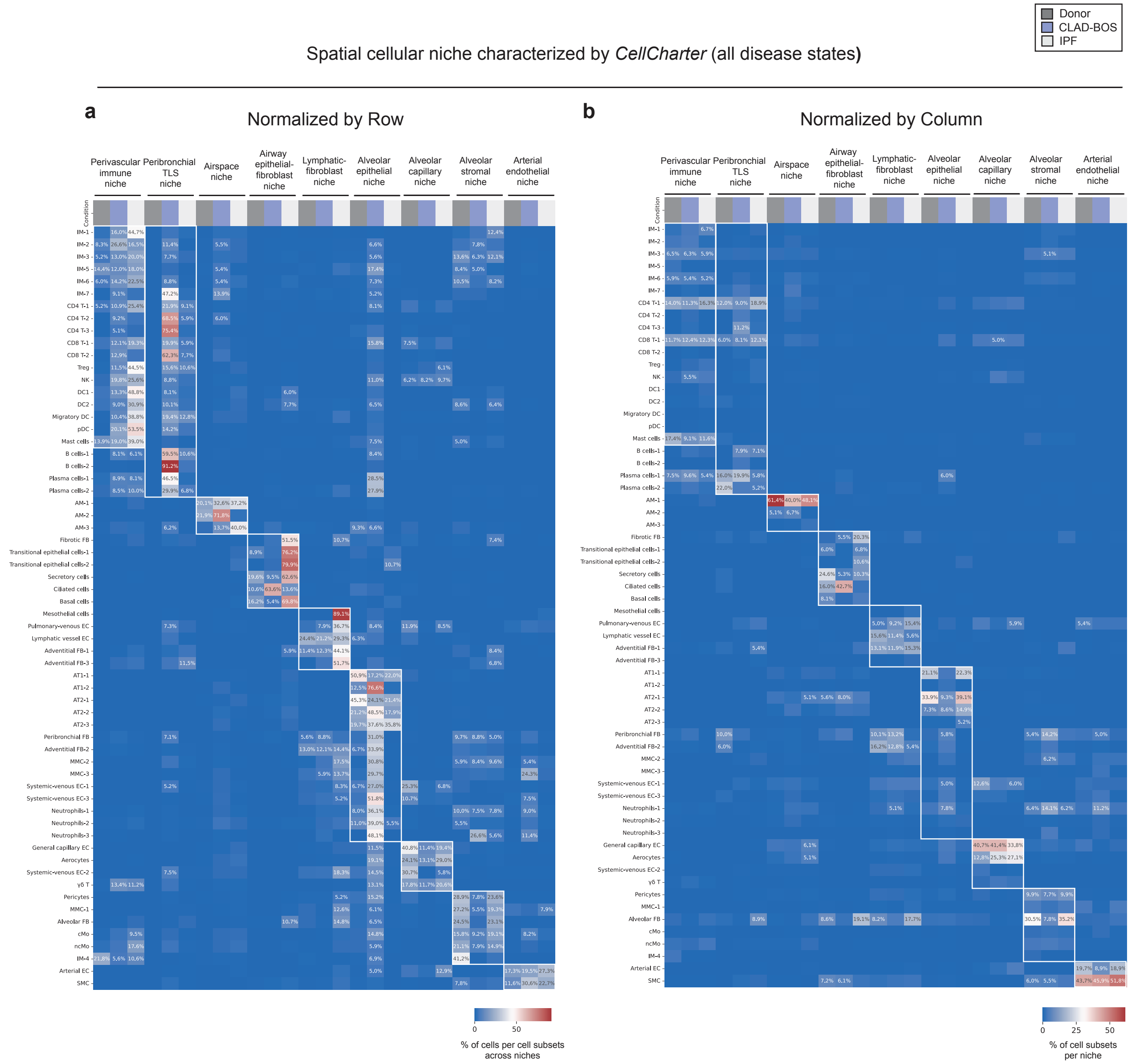


**Supplementary Figure S6. Spatial cellular niches in human lungs. a, b.** Composition of spatial cellular niches identified by *CellCharter* (a. normalized by row, b. normalized by column).

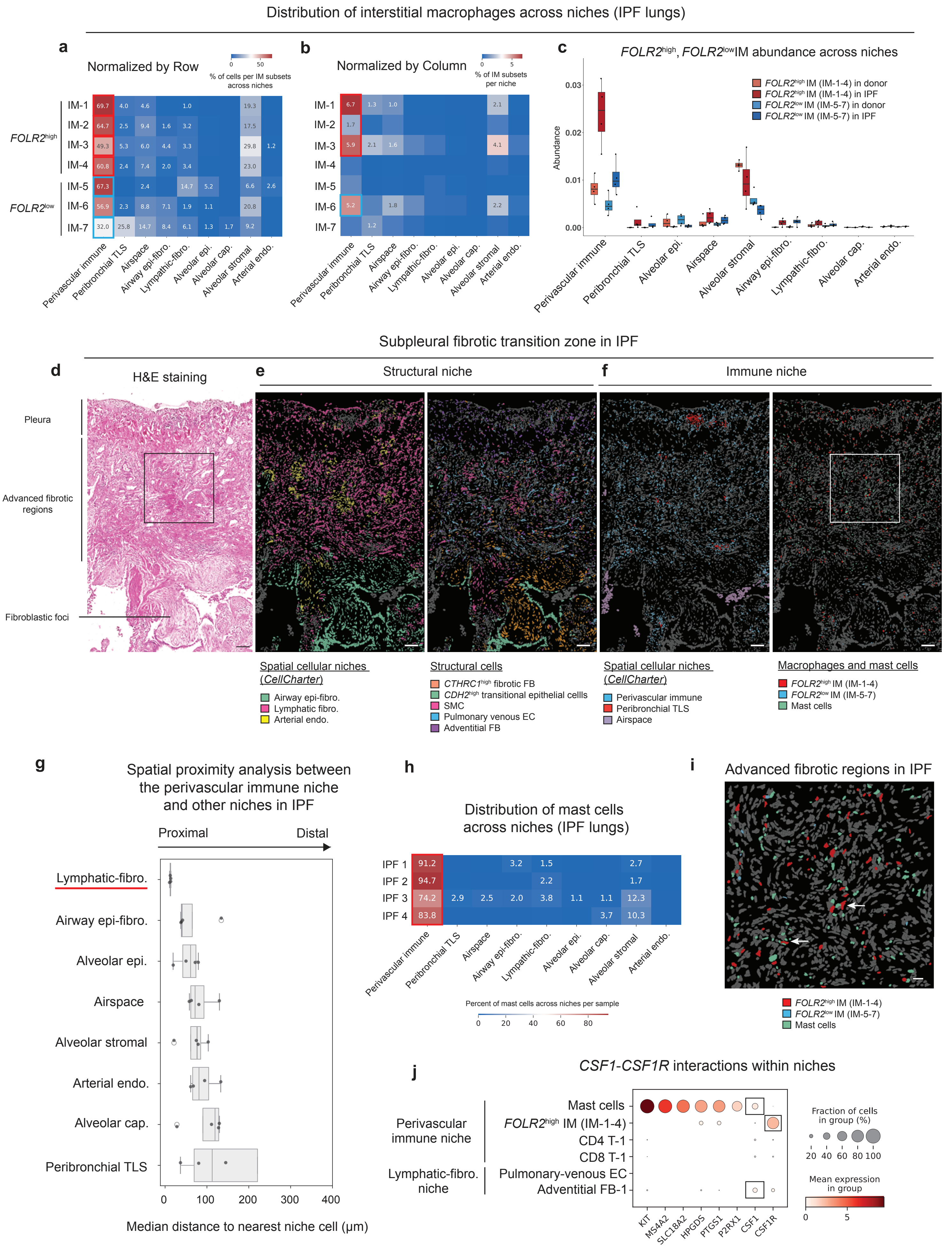

Distribution of interstitial macrophages across niches (IPF lungs)
a
Normalized by Row
% of cells per IM subsets across niches
FOLR2high
FOLR2low
IM-1
IM-2
IM-3
IM-4
IM-5
IM-6
IM-7
b
Normalized by Column
% of IM subsets per niche
c
FOLR2high, FOLR2low IM abundance across niches
FOLR2high IM (IM-1-4) in donor
FOLR2high IM (IM-1-4) in IPF
FOLR2low IM (IM-5-7) in donor
FOLR2low IM (IM-5-7) in IPF
Abundance
Perivascular immune
Peribronchial TLS
Airspace
Airway epi-fibro.
Lymphatic-fibro.
Alveolar epi.
Alveolar cap.
Alveolar stromal
Arterial endo.
Subpleural fibrotic transition zone in IPF
d
H&E staining
Pleura
Advanced fibrotic regions
Fibroblastic foci
e
Structural niche
f
Immune niche
Spatial cellular niches (CellCharter)
Airway epi-fibro.
Lymphatic fibro.
Arterial endo.
Structural cells
CTHRC1high fibrotic FB
CDH2high transitional epithelial cellls
SMC
Pulmonary venous EC
Adventitial FB
Spatial cellular niches (CellCharter)
Perivascular immune
Peribronchial TLS
Airspace
Macrophages and mast cells
FOLR2high IM (IM-1-4)
FOLR2low IM (IM-5-7)
Mast cells
g
Spatial proximity analysis between the perivascular immune niche and other niches in IPF
Proximal
Distal
Lymphatic-fibro.
Airway epi-fibro.
Alveolar epi.
Airspace
Alveolar stromal
Arterial endo.
Alveolar cap.
Peribronchial TLS
Median distance to nearest niche cell (μm)
h
Distribution of mast cells across niches (IPF lungs)
IPF 1
IPF 2
IPF 3
IPF 4
Percent of mast cells across niches per sample
i
Advanced fibrotic regions in IPF
FOLR2high IM (IM-1-4)
FOLR2low IM (IM-5-7)
Mast cells
j
CSF1-CSF1R interactions within niches
Perivascular immune niche
Lymphatic-fibro. niche
Mast cells
FOLR2high IM (IM-1-4)
CD4 T-1
CD8 T-1
Pulmonary-venous EC
Adventitial FB-1
KIT
MS4A2
SLC18A2
HPGDS
PTGS1
P2RX1
CSF1
CSF1R
Fraction of cells in group (%)
Mean expression in group

**Supplementary Figure S7. *FOLR2*$^{high}$ interstitial macrophages localize to specific niches in IPF lungs.** **a.** Distribution of interstitial macrophages across niches in IPF lungs (normalized by row). **b.** Distribution of interstitial macrophages across niches in IPF lungs (normalized by column). The heatmap displays the proportion of cells in each interstitial macrophage subset across niches, calculated by pooling cells from all samples. Values representing less than 1% are not shown. **c.** Relative abundance of *FOLR2*$^{high}$ and *FOLR2*$^{low}$ interstitial macrophages across niches, comparing donor to IPF lungs. Box plots represent the median and interquartile range. **d-f.** Representative hematoxylin and eosin-stained (H&E) sections and matching single-cell spatial transcriptomics images identifying the subpleural fibrotic transition zone in IPF. Box indicates the region of interest shown in Suppl. Figure S7i. **d.** H&E staining shows pulmonary pathologist-annotated fibroblastic foci, advanced fibrotic regions (defined by the accumulation of smooth muscle cells), and pleural regions. **e.** Distribution of structural niches and their constituent cell types. **f.** Distribution of immune niches and their constituent cell types. Selected cell types are highlighted in each panel. Scale bars 100 µm. **g.** Spatial proximity analysis between the perivascular immune niche and other niches in IPF lungs. Box plots showing the median absolute distance (µm) from edge cells within the perivascular niche to the nearest cells of other niches (n = 4). **h.** Distribution of mast cells across niches in IPF lungs. Heatmap displays the proportion of  mast cells across niches per sample (n = 4). **i.** Representative image showing the spatial proximity between *FOLR2*$^{high}$ interstitial macrophages and mast cells within the niche. Arrows indicate the co-localization of these two cell types. Scale bar 25 µm. **j.** Dot plot showing expression of genes distinguishing mast cells from other cell types within the perivascular immune and lymphatic-fibroblast niches. Only cell types constituting 10% or more of the niche are shown.

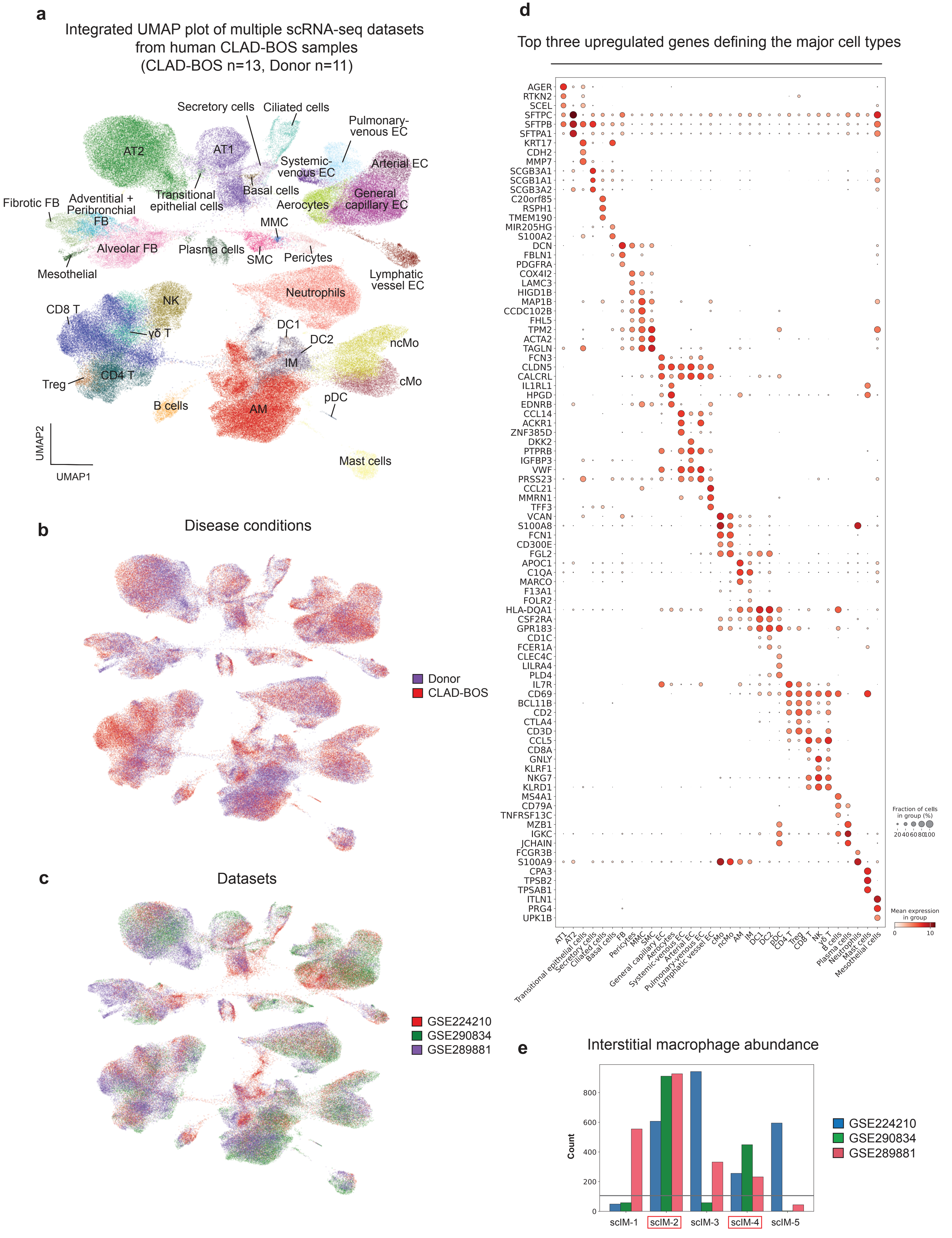
a
Integrated UMAP plot of multiple scRNA-seq datasets from human CLAD-BOS samples (CLAD-BOS n=13, Donor n=11)
Secretory cells
Ciliated cells
Pulmonary-venous EC
AT2
AT1
Systemic-venous EC
Arterial EC
Transitional epithelial cells
Basal cells
General capillary EC
Fibrotic FB
Adventitial + Peribronchial FB
Aerocytes
MMC
Alveolar FB
Plasma cells
SMC
Pericytes
Mesothelial
Lymphatic vessel EC
Neutrophils
NK
CD8 T
γδ T
DC1
DC2
ncMo
IM
Treg
CD4 T
cMo
pDC
B cells
AM
Mast cells
UMAP2
UMAP1
b
Disease conditions
Donor
CLAD-BOS
c
Datasets
GSE224210
GSE290834
GSE289881
d
Top three upregulated genes defining the major cell types
AGER
RTKN2
SCEL
SFTPC
SFTPB
SFTPA1
KRT17
CDH2
MMP7
SCGB3A1
SCGB1A1
SCGB3A2
C20orf85
RSPH1
TMEM190
MIR205HG
S100A2
DCN
FBLN1
PDGFRA
COX4I2
LAMC3
HIGD1B
MAP1B
CCDC102B
FHL5
TPM2
ACTA2
TAGLN
FCN3
CLDN5
CALCRL
IL1RL1
HPGD
EDNRB
CCL14
ACKR1
ZNF385D
DKK2
PTPRB
IGFBP3
VWF
PRSS23
CCL21
MMRN1
TFF3
VCAN
S100A8
FCN1
CD300E
FGL2
APOC1
C1QA
MARCO
F13A1
FOLR2
HLA-DQA1
CSF2RA
GPR183
CD1C
FCER1A
CLEC4C
LILRA4
PLD4
IL7R
CD69
BCL11B
CD2
CTLA4
CD3D
CCL5
CD8A
GNLY
KLRF1
NKG7
KLRD1
MS4A1
CD79A
TNFRSF13C
MZB1
IGKC
JCHAIN
FCGR3B
S100A9
CPA3
TPSB2
TPSAB1
ITLN1
PRG4
UPK1B
AT1
AT2
Transitional epithelial cells
Secretory cells
Ciliated cells
Basal cells
FB
Pericytes
MMC
SMC
General capillary EC
Aerocytes
Systemic-venous EC
Arterial EC
Pulmonary-venous EC
Lymphatic vessel EC
cMo
ncMo
AM
IM
DC1
DC2
pDC
CD4 T
Treg
CD8 T
NK
γδ T
B cells
Plasma cells
Neutrophils
Mast cells
Mesothelial cells
Fraction of cells in group (%)
20 40 60 80 100
Mean expression in group
0
10
e
Interstitial macrophage abundance
Count
0
200
400
600
800
scIM-1
scIM-2
scIM-3
scIM-4
scIM-5
GSE224210
GSE290834
GSE289881

**Supplementary Figure S8. Integrated scRNA-seq analysis of multiple published CLAD-BOS datasets. a.** UMAP visualization of the integrated scRNA-seq analysis across three independent, publicly available CLAD-BOS datasets (GSE224210, GSE290834, and GSE289881; total: CLAD-BOS, n = 13; Donor, n = 11). AM = alveolar macrophages; AT1 = alveolar epithelial type I cells; AT2 = alveolar epithelial type II cells; CD4 T = CD4 T cells; CD8 T = CD8 T cells; cMo = classical monocytes; DC = dendritic cells; EC = endothelial cells; FB = fibroblasts; γδ T = gamma delta T cells; IM = interstitial macrophages; MMC = microvascular mural cells; ncMo = non-classical monocytes; NK = natural killer cells; pDC = plasmacytoid dendritic cells; SMC = smooth muscle cells; Treg = regulatory T cells. **b.** UMAP plot colored by disease conditions. **c.** UMAP plot colored by dataset origin. **d.** Dot plot showing the top three marker genes for each major cell type. **e.** Abundance of interstitial macrophage clusters (scIM-1-5) across individual datasets. Only scIM-2 and scIM-4 were consistently identified across all analyzed datasets.

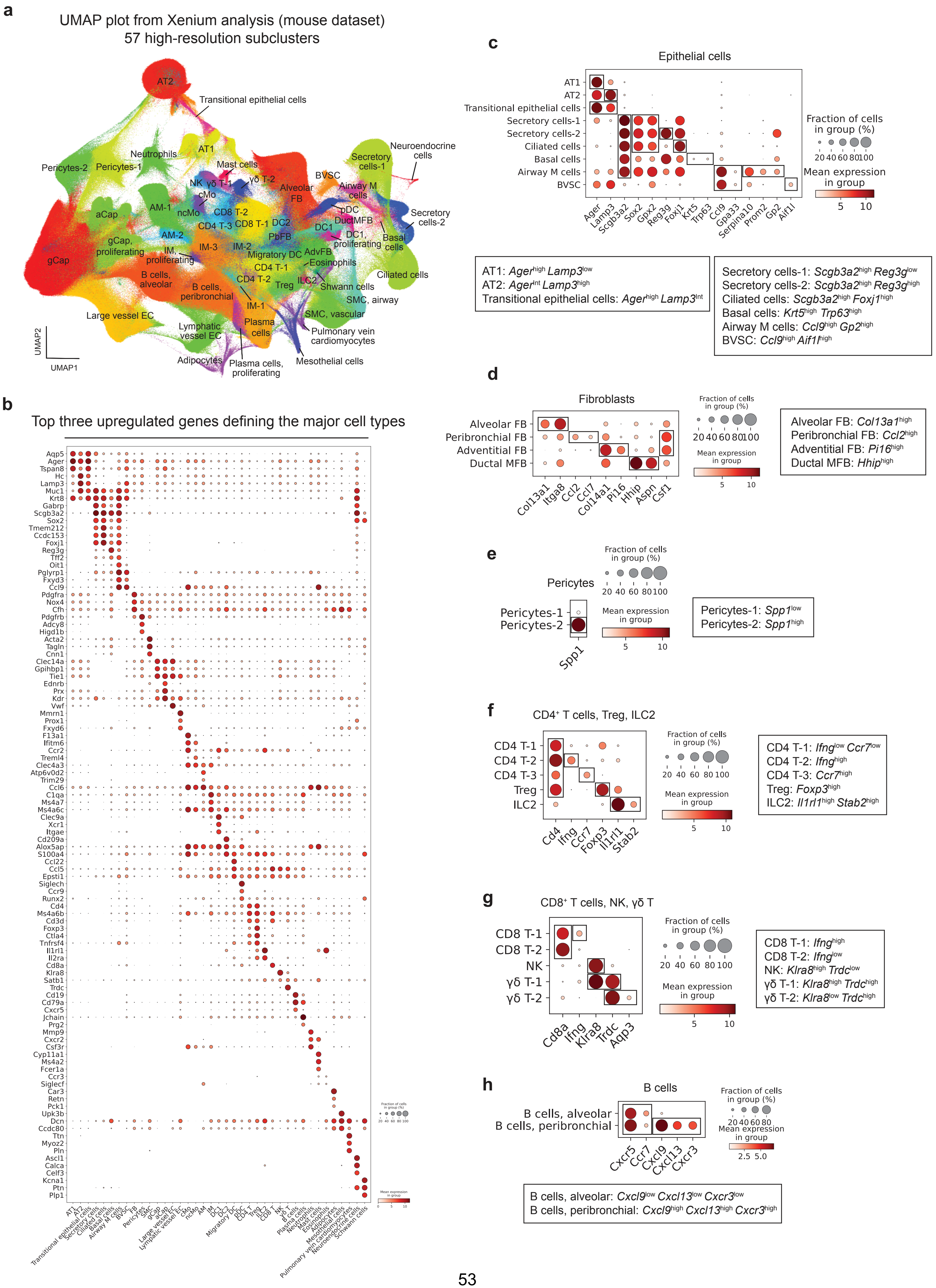
a
UMAP plot from Xenium analysis (mouse dataset)
57 high-resolution subclusters
AT2
Transitional epithelial cells
AT1
Neutrophils
Pericytes-2
Pericytes-1
Mast cells
NK γδ T-1
γδ T-2
cMo
ncMo
AM-1
aCap
CD8 T-2
CD4 T-3
CD8 T-1
DC2
AM-2
PbFB
gCap, proliferating
IM, proliferating
IM-3
IM-2
Migratory DC
AdvFB
gCap
CD4 T-1
Eosinophils
B cells, alveolar
B cells, peribronchial
CD4 T-2
Treg
ILC2
Shwann cells
IM-1
Large vessel EC
Lymphatic vessel EC
Plasma cells
Adipocytes
Plasma cells, proliferating
Mesothelial cells
Pulmonary vein cardiomyocytes
SMC, vascular
SMC, airway
Ciliated cells
Basal cells
DC1, proliferating
DC1
DuctMFB
pDC
Alveolar FB
Airway M cells
BVSC
Secretory cells-1
Neuroendocrine cells
Secretory cells-2
UMAP2
UMAP1
b
Top three upregulated genes defining the major cell types
c
Epithelial cells
Fraction of cells in group (%)
Mean expression in group
AT1: Ager^high Lamp3^low
AT2: Ager^int Lamp3^high
Transitional epithelial cells: Ager^high Lamp3^int
Secretory cells-1: Scgb3a2^high Reg3g^low
Secretory cells-2: Scgb3a2^high Reg3g^high
Ciliated cells: Scgb3a2^high Foxj1^high
Basal cells: Krt5^high Trp63^high
Airway M cells: Ccl9^high Gp2^high
BVSC: Ccl9^high Aif1l^high
d
Fibroblasts
Alveolar FB: Col13a1^high
Peribronchial FB: Ccl2^high
Adventitial FB: Pi16^high
Ductal MFB: Hhip^high
e
Pericytes
Pericytes-1: Spp1^low
Pericytes-2: Spp1^high
f
CD4+ T cells, Treg, ILC2
CD4 T-1: Ifng^low Ccr7^low
CD4 T-2: Ifng^high
CD4 T-3: Ccr7^high
Treg: Foxp3^high
ILC2: Il1rl1^high Stab2^high
g
CD8+ T cells, NK, γδ T
CD8 T-1: Ifng^high
CD8 T-2: Ifng^low
NK: Klra8^high Trdc^low
γδ T-1: Klra8^high Trdc^high
γδ T-2: Klra8^low Trdc^high
h
B cells
B cells, alveolar: Cxcl9^low Cxcl13^low Cxcr3^low
B cells, peribronchial: Cxcl9^high Cxcl13^high Cxcr3^high

**Supplementary Figure S9. Single-cell spatial transcriptomics resolves cell types in the mouse lung. a.** UMAP visualization of single-cell spatial transcriptomic data from murine experiments (total number of cells = 3,502,604 cells, 479 genes, 57 high-resolution subclusters). Lung allografts and syngeneic grafts from three experimental groups (n = 3 per group; total n = 9) were included in this UMAP. aCap = alveolar capillary endothelial cells; AM = alveolar macrophages; AT1 = alveolar epithelial type I cells; AT2 = alveolar epithelial type II cells; BVSC = bronchovascular bundle sheath cells; CD4 T = CD4 T cells; CD8 T = CD8 T cells; cMo = classical monocytes; DC = dendritic cells; EC = endothelial cells; FB = fibroblasts; gCap = general capillary endothelial cells; γδ T = gamma delta T cells; ILC2 = group 2 innate lymphoid cells; IM = interstitial macrophages; MFB = myofibroblasts; ncMo = non-classical monocytes; NK = natural killer cells; pDC = plasmacytoid dendritic cells; SMC = smooth muscle cells; Treg = regulatory T cells. **b.** Dot plot showing the top three marker genes for each major cell type. **c.** Dot plot showing selected genes used to resolve epithelial cell types. **d.** Dot plot showing selected genes used to resolve fibroblast subtypes. **e.** Dot plot showing selected genes used to resolve pericyte subtypes. **f.** Dot plot showing selected genes used to resolve CD4 T cells, Treg, and ILC2. **g.** Dot plot showing selected genes used to resolve CD8 T cells, NK cells, and γδ T cells. **h.** Dot plot showing selected genes used to resolve B cell subsets.

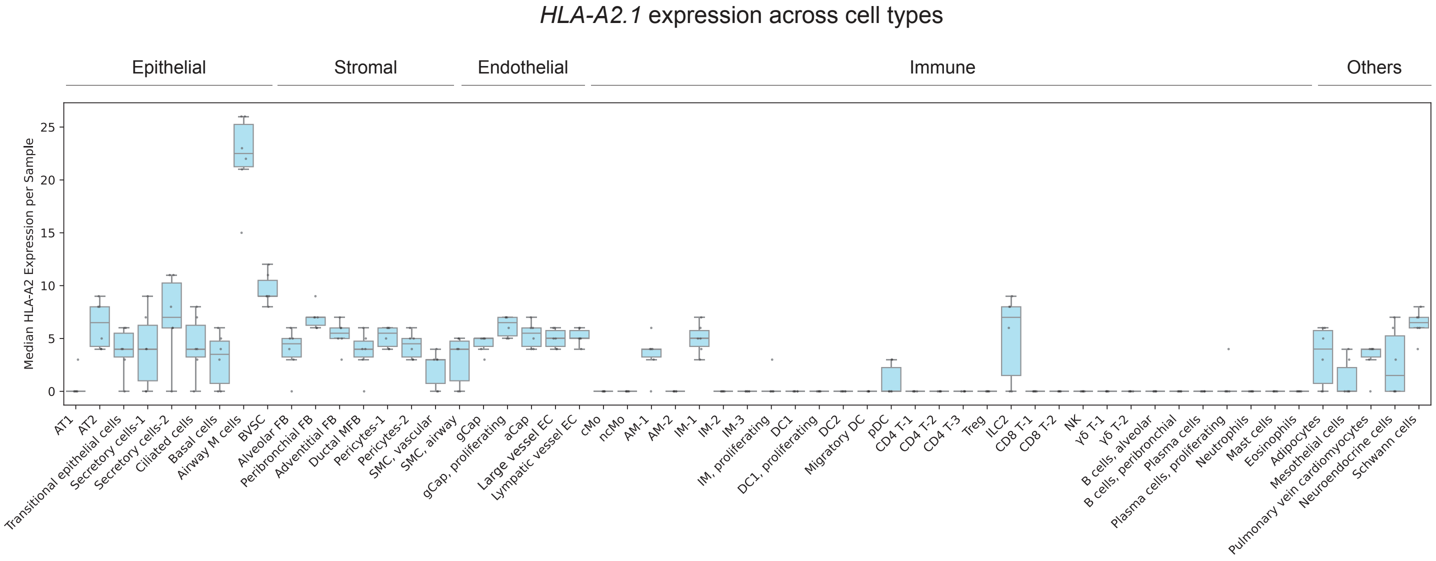


**Supplementary Figure S10. HLA-A2.1 lineage tracing resolves donor- and recipient-derived cells in murine CLAD-BOS.** HLA-A2.1 expression across all cell types. Samples from single-mismatch lung transplantations (n = 6) were included in this analysis. Box plots represent the median and interquartile range.

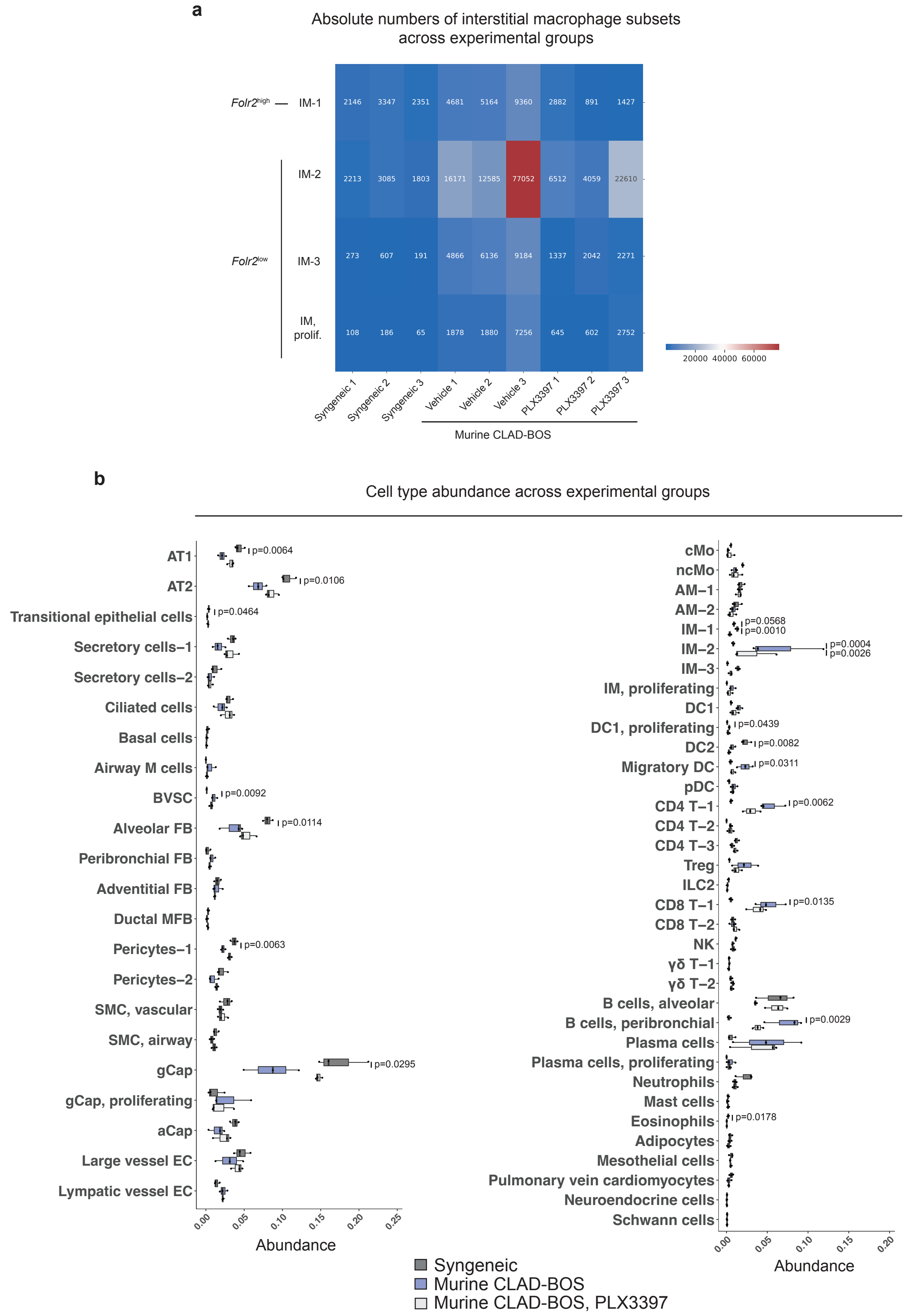


**Supplementary Figure S11. Cell type abundance between syngeneic and allogeneic transplant lungs using per-sample fractions. a.** Absolute numbers of interstitial macrophages across experimental conditions. *Folr2*$^{high}$ interstitial macrophages = IM-1; *Folr2*$^{low}$ interstitial macrophages = IM-2, 3, prolif. **b.** Differential abundance analysis based on per-sample fractions. Box plots represent the median and interquartile range. Statistical significance was assessed using pairwise t-tests with Bonferroni correction for multiple comparisons.

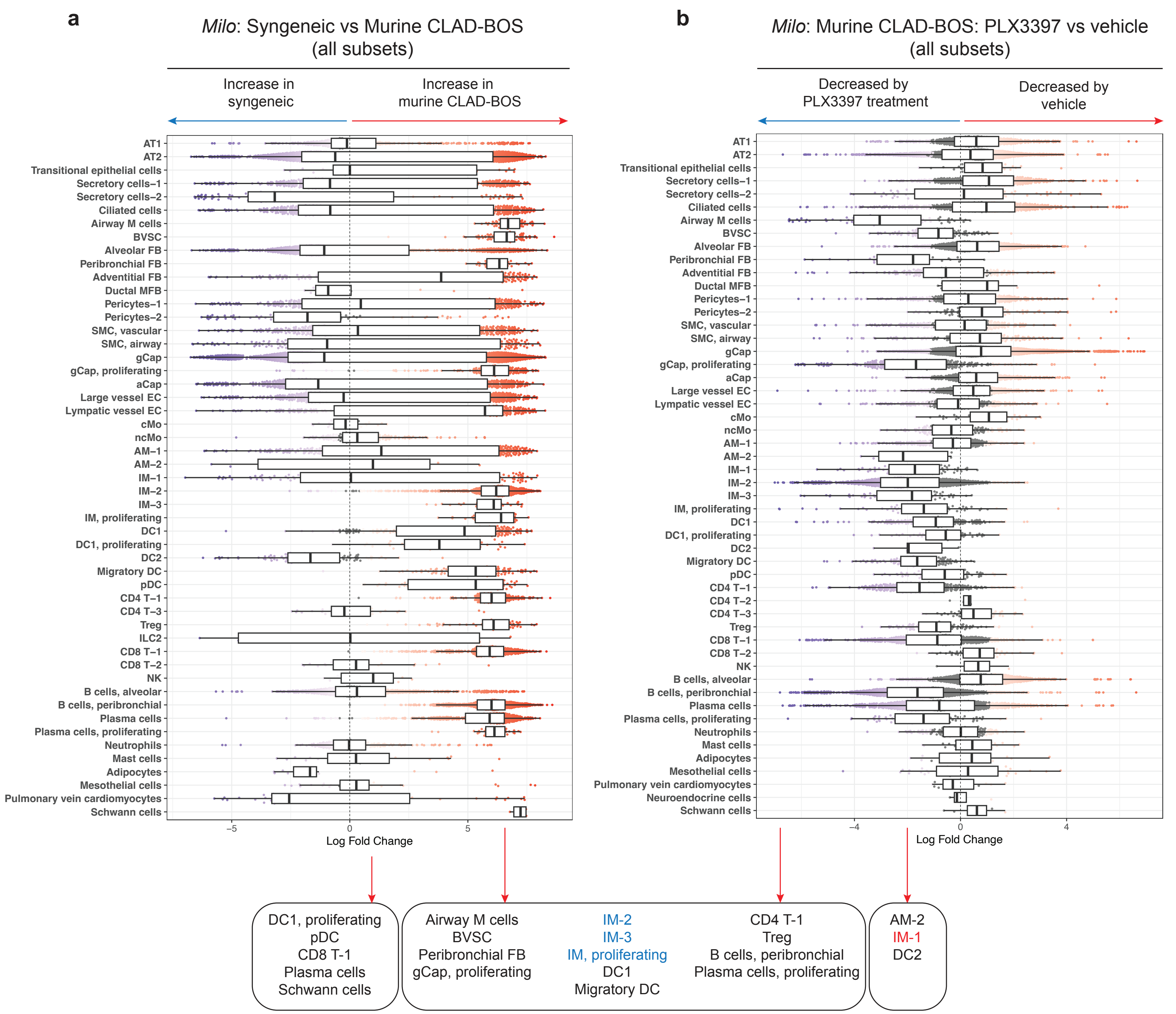


**Supplementary Figure S12. Cell type abundance between syngeneic and allogeneic transplant lungs using *Milo*.** Differential abundance analysis using *Milo* showing all cell types in murine CLAD-BOS lungs relative to syngeneic grafts with or without PLX3397 treatment. Box plots represent the median and interquartile range. Cell populations were defined as significantly changed if at least four neighborhoods were identified, and both the upper and lower quartiles of Log-fold change were skewed to one side relative to zero.

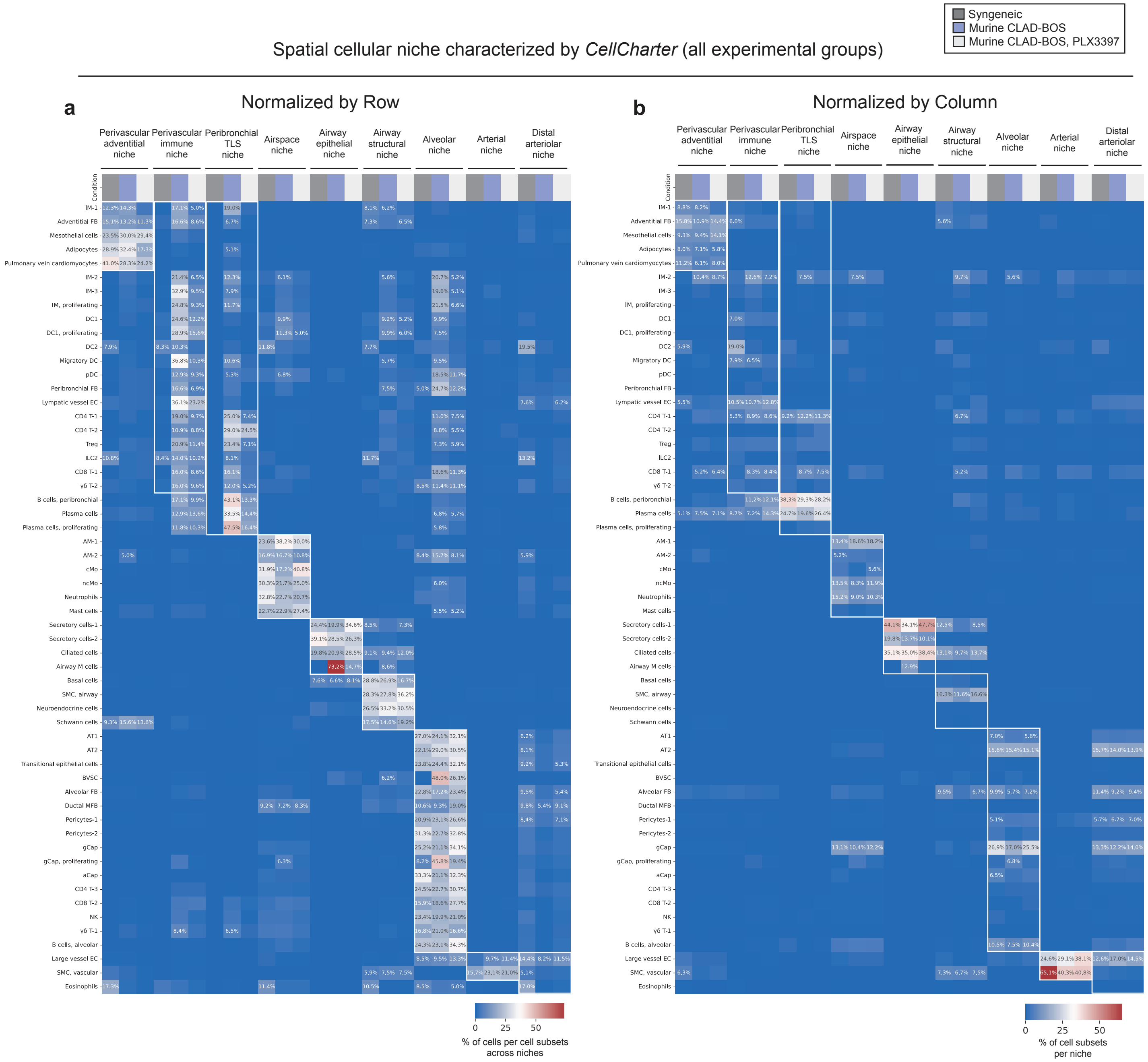


**Supplementary Figure S13. Spatial cellular niches in mouse lungs. a, b.** Composition of spatial cellular niches identified by *CellCharter* (a. normalized by row, b. normalized by column).

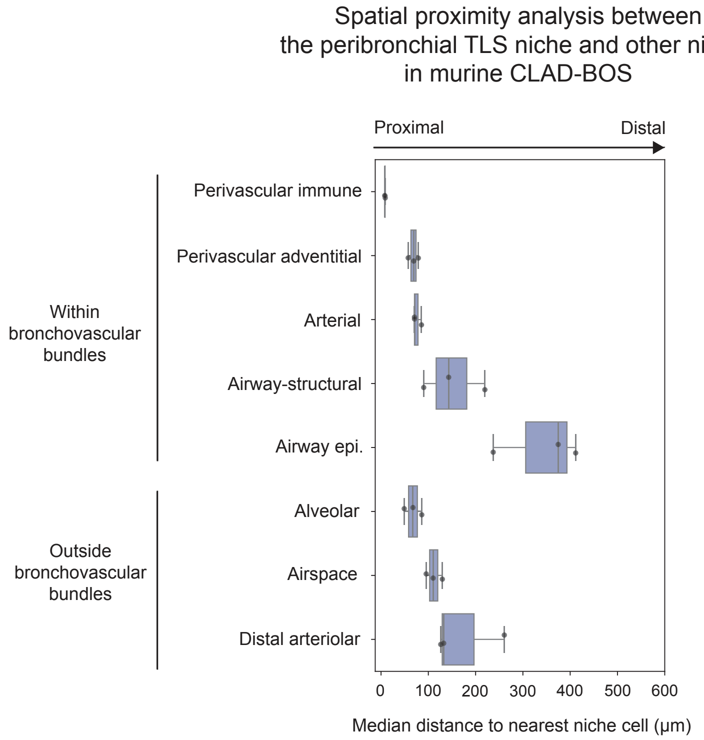


**Supplementary Figure S14. Spatial proximity analysis between the peribronchial TLS niche and other niches in murine CLAD-BOS lungs.** Box plots showing the median absolute distance (μm) from edge cells within the peribronchial TLS niche to the nearest cells of other niches (n = 3).

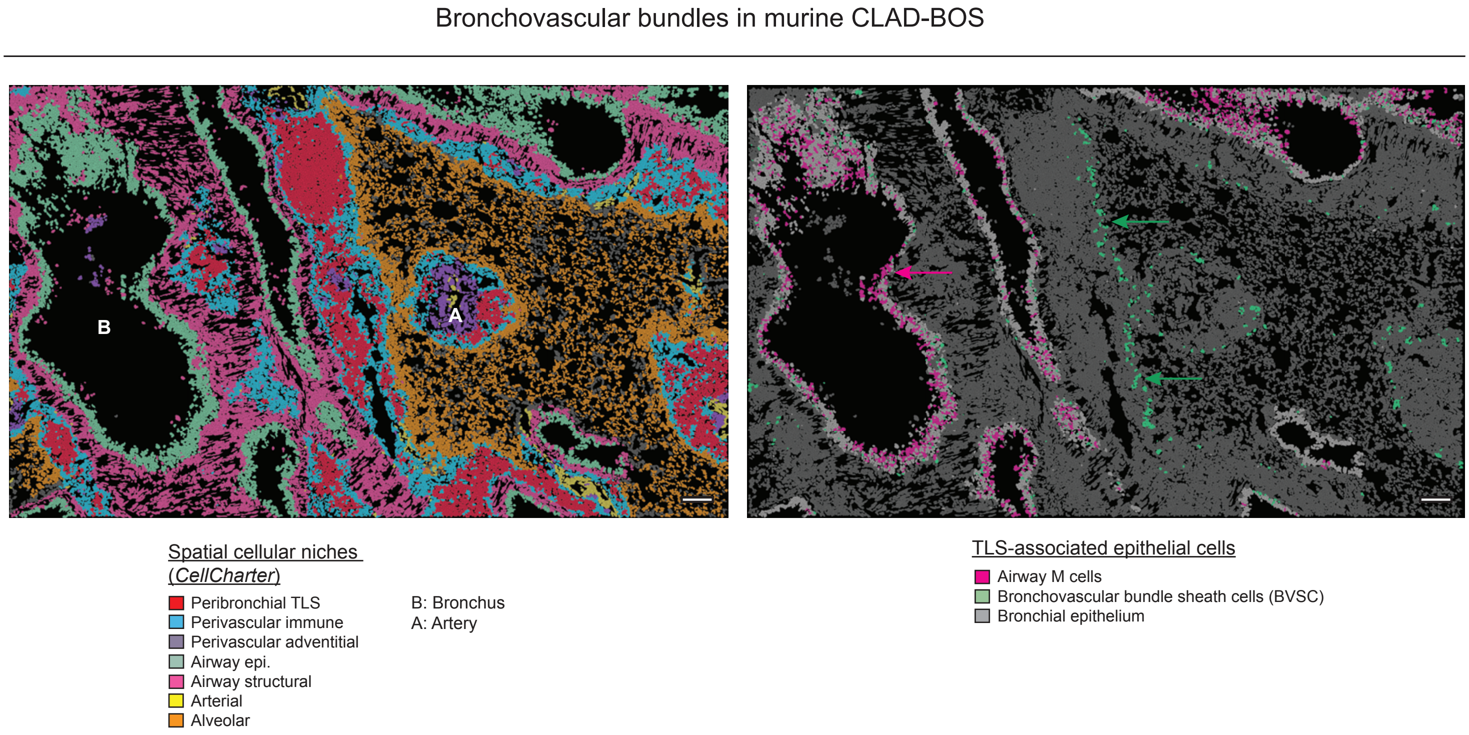


**Supplementary Figure S15. Single-cell spatial transcriptomics resolves distinct epithelial cell states associated with TLS formation in murine CLAD-BOS.** Representative images highlighting spatial cellular niches and the distribution of TLS-associated epithelial cells (airway M cells, BVSC) in murine CLAD-BOS. Selected cell types are highlighted in each panel. Scale bar 100 μm.

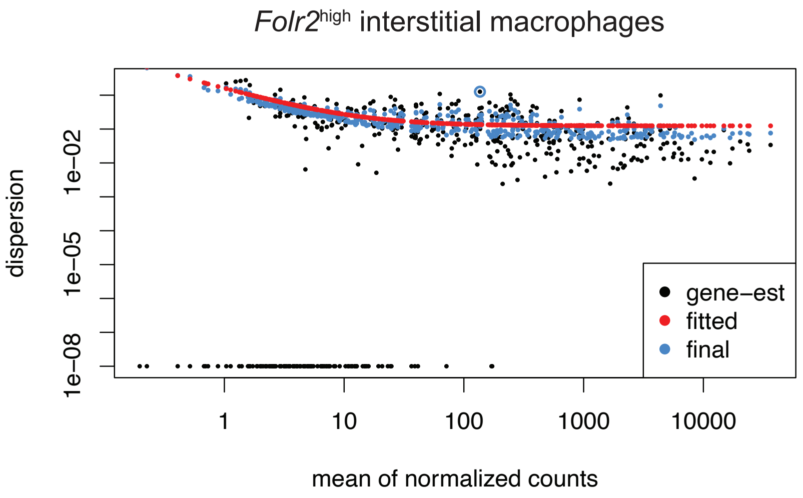


**Supplementary Figure S16. Dispersion estimation for differential gene expression of *Folr2*high interstitial macrophages in murine CLAD-BOS.** Dispersion estimation of transcripts in *Folr2*high interstitial macrophages. Mean-variance relationship of transcript data conforms to the negative binomial distribution. The red line indicates the fitted dispersion trend used for normalization.

**Supplementary Table legends**

**Supplementary Table S1. Patient characteristics**

**Supplementary Table S2. Gene list for the Xenium human panel**

**Supplementary Table S3. Gene list for the Xenium mouse panel**

**Supplementary Table S4. DEGs for scIM in integrated CLAD-BOS scRNA-seq**

**Supplementary Table S5. DEGs in *Folr2*$^{high}$ interstitial macrophages comparing peribronchial TLS niche and perivascular immune niche**
