## Supplementary material for "Interstitial macrophages drive chronic lung allograft dysfunction": Suppl. Table S1

**Supplementary Table S1. Patient characteristics**

|  | **Sex** | **Age** | **Primary lung diseases** |
| --- | --- | --- | --- |
| Donor-1 | Male | 36 | N/A |
| Donor-2 | Female | 32 | N/A |
| Donor-3 | Female | 28 | N/A |
| Donor-4 | Female | 35 | N/A |
| CLAD-BOS 1 | Male | 28 | Cystic fibrosis |
| CLAD-BOS 2 | Female | 43 | Cystic fibrosis |
| CLAD-BOS 3 | Male | 64 | COPD |
| IPF-1 | Male | 60 | IPF |
| IPF-2 | Male | 72 | IPF |
| IPF-3 | Male | 74 | IPF |
| IPF-4 | Female | 64 | IPF |

COPD = Chronic obstructive pulmonary diseases
